# *In vivo* interrogation of transcriptional and epigenetic regulators of lung epithelial regeneration

**DOI:** 10.64898/2026.03.13.711474

**Authors:** Dawei Sun, Daisy A. Hoagland, Daniel Strebinger, Chenlei Hu, Benno C. Orr, Shahinoor Begum, Giovanni Marrero, Jackson A. Weir, Ruth A. Franklin, Fei Chen

## Abstract

Effective alveolar repair after viral lung injury requires precise coordination of alveolar type 2 cell (AT2) proliferation and differentiation to restore lung function. To uncover causal regulators of this process in the native tissue environment, we developed SAGE (<u>S</u>table <u>A</u>deno-Associated Virus <u>G</u>enomic Int<u>E</u>gration), an engineered AAV system that enables high-throughput *in vivo* genetic interrogation. SAGE supports both bulk phenotypic screening (SAGE-Perturb) and single-cell transcriptomic profiling (SAGE-Perturb-seq). Using this approach, we identified lysine acetyltransferase 8 (Kat8) as essential for epithelial repair following viral infection through the Non-Specific-Lethal (NSL) complex, and generated a time-resolved, high-resolution functional map of transcription factor knockouts during alveolar repair, revealing transcription factor dependences for distinct alveolar epithelial repair trajectories. This map further defined two independent AT2-derived transitional states: a reparative state, and a pathological state that is transcriptionally similar to the basaloid population observed in human pulmonary fibrosis. Disruption of transcription factors in the NF-*κ*B pathway prevented the emergence of the pathological transitional state, linking inflammation and maladaptive epithelial remodeling. SAGE represents a versatile platform for functional genomics *in vivo*, with applications extending across respiratory biology and disease.

## Introduction

The respiratory tract is continuously exposed to viruses and has evolved robust mechanisms to rapidly eliminate infection and restore tissue integrity. Within the alveolar regions of the lung, efficient repair is especially critical: impaired regeneration can lead to life-threatening conditions such as acute respiratory distress syndrome (ARDS), while incomplete or aberrant repair may drive the progression of chronic lung diseases, including pulmonary fibrosis. Alveolar type 2 epithelial cells (AT2s) serve as key epithelial progenitors that self-renew and differentiate into alveolar type 1 cells (AT1s) during alveolar regeneration. This dynamic regenerative process is orchestrated by multiple factors, including extrinsic signaling cues^1–6^, transcriptional and epigenetic regulation^7–10^, cellular metabolism^11,12^, and mechanical forces^13^. Recent studies have identified transitional states, including damaged-associated transient progenitors (DATPs), also known as Krt8^+^ alveolar differentiation intermediates (ADIs) or pre-alveolar type-1 transitional cells (PATS), which arise as AT2 differentiate into AT1 during repair^6,14–16^. While these intermediates facilitate recovery after acute lung injury, their persistence during chronic inflammation has been proposed to contribute to fibrosis and cancer^17–21^. Despite this progress, a comprehensive understanding of how epithelial-intrinsic transcriptional and epigenetic programs integrate with extrinsic signals to govern AT2 fate, including quiescence, self-renewal, differentiation, or dysplastic transformation, remains incomplete.

Clustered regularly interspaced short repeats (CRISPR) gene editing technology allows large-scale genetic screens in diverse biological systems, including cell lines, organoids, and, more recently, *in vivo* models^22–31^. Furthermore, by combining CRISPR screening with single-cell and imaging-based readouts, transcriptomic and protein-level outputs can be utilized for high dimensional phenotyping of genetic perturbations within native tissue environments^32–34^. However, such a high-throughput *in vivo* genetic screening platform has not yet been established for the mammalian lung.

Here, we developed SAGE, a genome-integrating adeno-associated virus (AAV) that delivers guide RNA (gRNA) libraries to the lung epithelium, enabling bulk phenotypic screens and single-cell Peturb-seq profiling directly from mouse lungs *in vivo*. Applying this platform, we screened a curated library of 170 chromatin modulators in a mouse model of influenza A virus (IAV)-induced lung injury and identified *Kat8*, along with other histone acetylation-related genes, as essential regulators of AT2-mediated regeneration. We further demonstrated that the function of *Kat8* is mediated through the non-specific lethal (NSL) complex and that disruption of NSL complex function induced stress responses in AT2s, leading to senescence-like phenotypes and expression of profibrotic factors. To better understand the requirements of transcription factors (TFs) in this repair process, we constructed a time-resolved SAGE-Perturb-seq atlas encompassing 118 TF perturbations. This atlas revealed four distinct meta-functional groups of TFs that cooperatively shape, bias, or destabilize epithelial regeneration trajectories following injury. It also resolves two distinct intermediate cell states that recapitulate the reparative and pathological transitional populations recently observed in diseased human lungs^17,19,21^. For the first time, we were able to identify the genetic drivers of these transitional states and demonstrated that elevated inflammatory signaling, including NF-*κ*B, fuels the emergence of the pathological population. Together, these findings establish SAGE-Perturb and SAGE-Perturb-seq as a scalable *in vivo* screening framework broadly applicable to lung injury models and potentially other regenerating tissues, enabling systematic dissection of the genetic circuits that orchestrate tissue repair.

## Results

### An AAV-Sleeping Beauty hybrid system provides efficient transduction in the mouse lung

AAVs are among the most widely used vectors for *in vivo* delivery due to their broad tissue tropism, low immunogenicity, and long-term expression in non-dividing cells. These features have made them attractive for gene therapy and single-gene functional studies across organ systems^35^. However, two major limitations have restricted their utility for high-throughput *in vivo* genetic screening. First, AAV tropism varies across serotypes and often fails to achieve efficient, cell-type-specific transduction at low multiplicity of infection, a prerequisite for pooled perturbation approaches. Second, AAV genomes typically remain episomal^36^ and are progressively diluted during cell division, preventing long-term tracking of perturbations by sequencing single gRNAs in proliferating cells, a key requirement for studying regeneration in mitotic tissues such as the lung epithelium.

To overcome these challenges, we sought to develop a stable, low-multiplicity, genome-integrating AAV platform that would enable pooled *in vivo* screening in the regenerating lung. We began by identifying an efficient AAV serotype for *in vivo* AT2 transduction. We first compared AAV2, AAV5, AAV6 and AAV9 serotypes delivered intratracheally (i.t.). AAV9 demonstrated both high AT2 selectivity and high *in vivo* AT2 transduction efficiency (∼20%, N = 4, **Fig. 1A**), consistent with previous reports^1,37^.

**Fig. 1.**
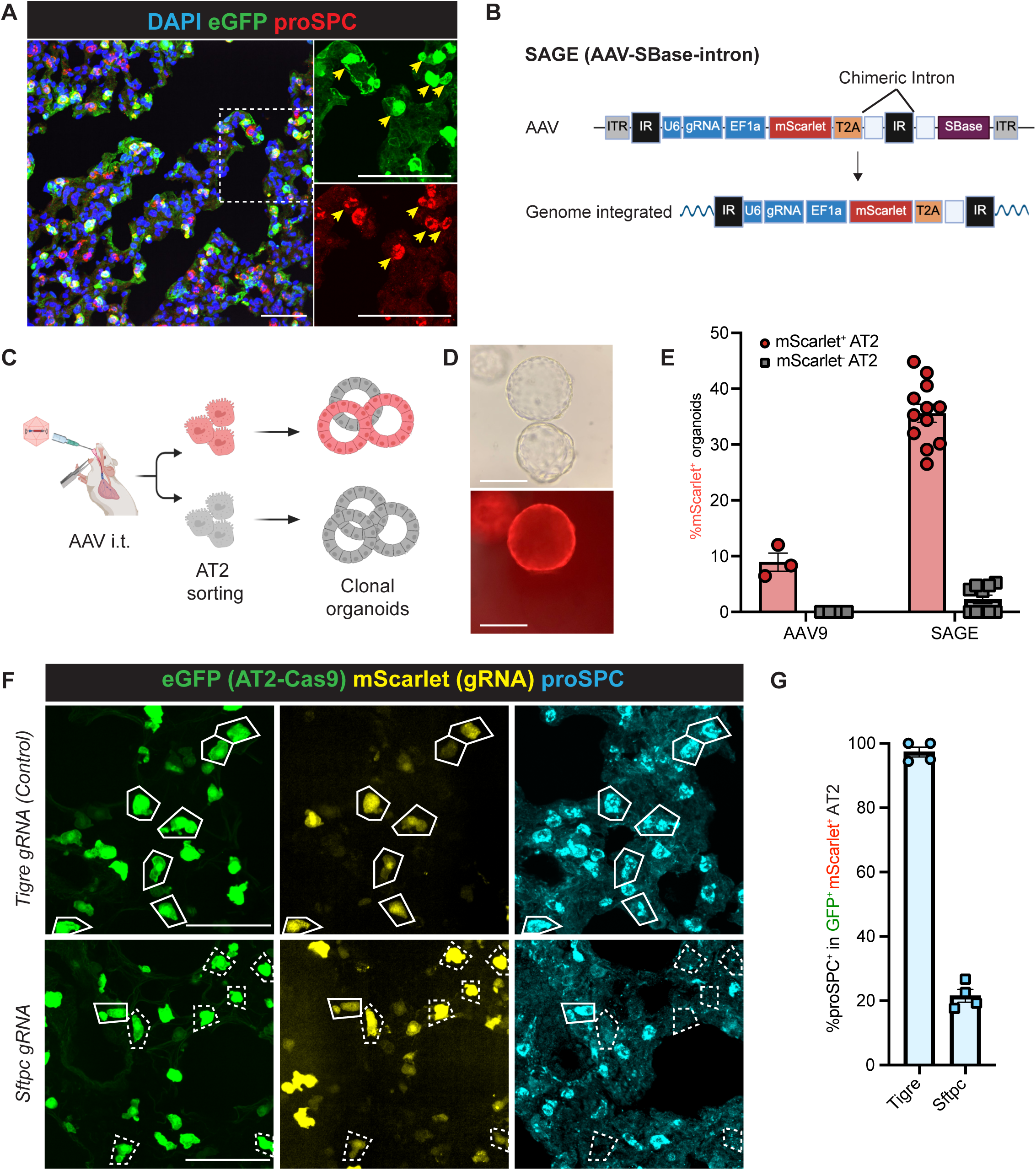
Establishment of an AAV-based *in vivo* genetic screening platform in mouse lung. **(A).** Immunofluorescence (IF) images of *in vivo* AT2 transduction using AAV9. Green: eGFP; Red: proSPC. Yellow arrows indicate the AT2s (proSPC^+^) that are infected by AAV9-eGFP. Scale bars, 50 μm. **(B).** Schematic of the SAGE (AAV-SBase-intron) system. **(C).** Experimental design for quantification of genome integration efficiency of engineered AAV systems. Lungs were harvested from mice seven days after i.t. administration. **(D).** Representative widefield images of the organoids grown from AT2s transduced by SAGE. mScarlet^+^ AT2s were harvested as CD45^-^Pdgfra^-^CD31^-^EpCAM^+^MHCII^+^CD104^-^. Scale bars, 200 μm. **(E).** Quantification for integration efficiency of AAV9 and SAGE using the AT2 clonal organoid assay. N = 3 and 12 mice were used for AAV9 and SAGE, respectively. Error bars: mean ± SEM. **(F).** IF images illustrating *Sftpc* gene knockout in AT2s. AT2-specific knockouts were achieved through *AT2-Cas9-eGFP* mice following tamoxifen treatment and subsequent SAGE gRNA delivery (detailed experimental scheme in Fig. S1C). A gRNA targeting the *Tigre* locus^93^, a genomic safe harbor in the mouse genome, was used as a control. Green: eGFP representing Cas9 expression; yellow: mScarlet, representing SAGE transduction; cyan, proSPC. White dashed lines indicate AT2s expressing Cas9 and gRNA that have lost proSPC expression. White solid lines indicate AT2s expressing Cas9 and gRNA that have retained proSPC expression. Scale bars, 50 μm. **(G).** Quantification of proSPC knockout effects in AT2s that are mScarlet^+^eGFP^+^. N = 4 mice were used for each condition. Error bars: mean ± SEM.

To achieve stable integration using an AAV vector, we leveraged the Sleeping Beauty (SB) transposon system, a cut and paste DNA transposition system, which is commonly used in vertebrate cells due to its high efficiency^38^. SB consists of a transposase enzyme (SBase) and flanking inverted terminal repeats (IRs), with SBase catalyzing excision of the transposon and insertion into genomic DNA at TA dinucleotides. We tested two hybrid AAV-SB systems, both of which encoded an mScarlet reporter and the SBase, but differed in their regulation of transposase expression: (1) AAV-SBase^27,28^ (**fig. S1A**), where SBase catalyses genome integration after AAV internalization but remains expressed after integration and (2) AAV-SBase-intron (**Fig. 1B**), where an SB IR is embedded within a chimeric intron, such that SBase expression is lost after genome integration, thereby preventing remobilization of the transgene.

To evaluate the genome integration efficiency of these two systems, we transduced AT2s *in vivo* by i.t. administration of AAVs and isolated the mScarlet^+^ transduced cells for a clonal organoid assay (**Fig. 1C**). For comparison, we also transduced AT2s *in vivo* with unmodified AAV9. In this assay, only AT2s that have undergone genome integration will give rise to organoids that are uniformly mScarlet positive. Both the AAV-SBase and the AAV-SBase-intron systems yielded markedly higher percentages of mScarlet^+^ organoids (28.2% and 35.6% respectively) in comparison to unmodified AAV9 (**Fig. 1, D and E and fig. S1B**), confirming their enhanced integration efficiency. Given that the AAV-SBase-intron design further stabilizes the transgene by eliminating SBase expression, we selected this design for subsequent experiments and refer to it as SAGE (<u>S</u>table <u>A</u>AV <u>G</u>enomic Int<u>E</u>gration).

### SAGE allows efficient *in vivo* gene editing in the mouse lung at low multiplicity of infection

To assess the suitability of SAGE for pooled screening in the lung, we first tested whether SAGE could achieve efficient gene editing *in vivo*. We delivered *Sftpc*-targeting gRNAs to *Sftpc-CreER^+/–^; LSL-Cas9-2A-eGFP^+/–^* mice (hereafter AT2-Cas9-eGFP). These mice allow tamoxifen-induced recombination to activate eGFP and Cas9 expression, which upon successful gRNA delivery results in the knockout of *Sftpc*, a surfactant protein gene expressed in AT2s. Two weeks after SAGE delivery, we observed efficient knockout of *Sftpc* at both the genomic and protein levels (78.5%, N = 4; **Fig. 1, F and G** and **fig.1, C, D and E**), confirming efficient gene-editing capability of the system *in vivo*.

Next, we quantified the rate of AAV multi-infection in AT2s by performing a dual-color co-transduction experiment using equal ratios of AAVs expressing eGFP and mCherry, delivered i.t. at increasing total viral genome (vg) doses (from 4 × 10^9^ to 1 × 10^11^). We quantified single- and dual-labeled AT2s by flow cytometry after 5 days. We observed a dose-dependent transduction of AT2s, with a low dual-infection rate (∼4%) at the highest dose tested (**fig. S2A**), indicating that SAGE enables efficient delivery while favoring single-copy uptake. We therefore selected 1 × 10^11^ vg as the working viral titer for subsequent experiments. Together, these established SAGE as a suitable platform for *in vivo* genetic screening applications.

### SAGE enables *in vivo* pooled genetic screens in the regenerating lung epithelium

We next evaluated the performance of SAGE for *in vivo* genetic screens during lung repair. We used a well-established IAV-induced lung injury model^5,39^, which causes acute epithelial damage and triggers robust AT2 proliferation and differentiation. This system provides a defined temporal window to systematically interrogate the genetic regulators of AT2-mediated repair. Importantly, it also recapitulates key features of viral lung injury and regeneration observed in humans^40^. Moreover, injury models offer a critical advantage, as AT2s are largely quiescent under homeostatic conditions^1,2,41^, making it difficult to capture their proliferation and differentiation within a practical screening window.

We focused our screens on two major classes of intrinsic regulators of cell states and fates: chromatin modulators (CMs) and TFs^7–10,42–45^. For CMs, we assembled a library targeting 173 CMs, spanning major functional categories^26,46^, with an average of 3 gRNAs per gene. For TFs, we curated a library targeting 118 TFs that are enriched in AT2s, AT1s, or transitional states across lung development and various injury models^2,6,47^, using an average of 5 gRNAs per gene. Both libraries included multiple types of controls: non-targeting gRNAs with no binding sites in the genome, targeting controls directed at intergenic regions or safe harbor loci, and positive controls consisting of gRNAs targeting essential genes^22^ (**table S1**). We estimated that libraries of this size can easily be well represented with >500 AT2s/gRNA within a single mouse (**Methods**).

To identify CMs and TFs essential for epithelial repair after injury, CM and TF libraries were independently cloned into the SAGE system and administered i.t. to transduce AT2s. After a 12-14 day period to allow for genomic integration and gene knockout, mice were challenged with IAV. 28 days post-infection (28 dpi), a time point following viral clearance and alveolar repair^39^, we isolated AT2s from lung samples using an EpCAM^+^MHCII^+^ gating strategy^48^. We then quantified gRNA representation in the isolated AT2s via next generation sequencing (**Fig. 2A**).

**Fig. 2.**
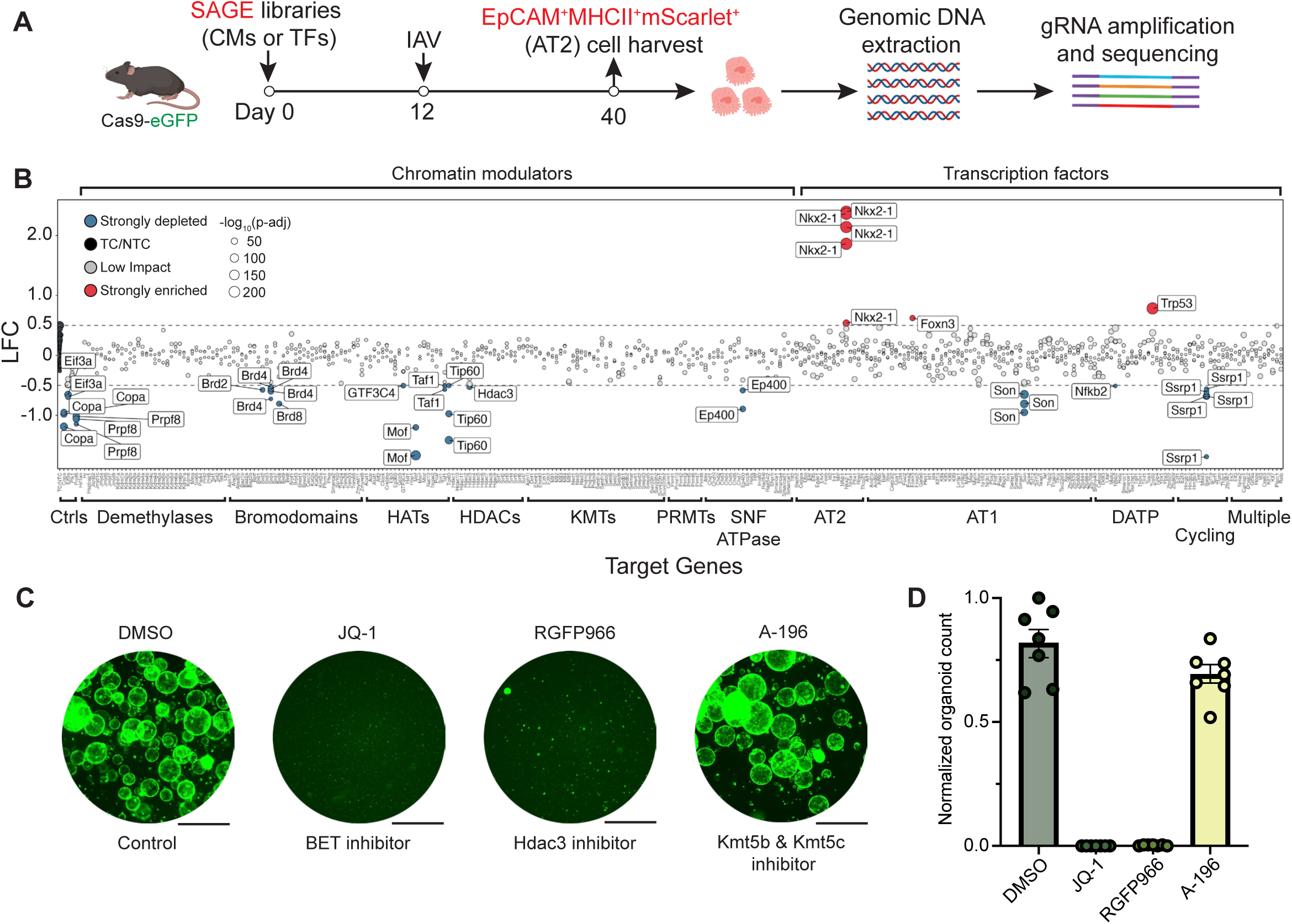
*In vivo* screens identified essential regulators of alveolar repair. **(A).** Schematic workflow for *in vivo* genetic screens in mouse lung using SAGE-Perturb. Constitutive Cas9-eGFP mice were i.t. treated with SAGE CM and TF libraries before infection with IAV. mScarlet positive AT2s (CD45^-^CD31^-^Pdgfra^-^EpCAM^+^CD104^-^MHCII^+^) were sorted 28 days after IAV infection, followed by DNA extraction, library amplification, and sequencing. N = 5 mice were used for each library. **(B).** Changes in gRNA abundance. Each circle represents a single gRNA, with gRNAs targeting the same gene aligned vertically. Circle size corresponds to −log_10_(adjusted p-value). Targeting controls (TC) and non-targeting controls (NTC), are shown as filled black circles. Strongly enriched and depleted gRNAs (defined by log_2_ fold change (LFC) > 0.5 or < −0.5 and adjusted p-value < 0.05) are color-coded in red and blue respectively. For CMs, categories are labeled below. HATs, histone acetyltransferases. HDACs, histone deacetylases. KMTs, histone lysine methyltransferases. PRMTs, protein arginine methyltransferases. For TFs, enriched cell populations from publicly available datasets are labeled below. Multiple, TFs enriched in more than one cell population as defined by scRNA-seq. **(C).** Representative images showing AT2 organoid growth under chromatin modulator inhibitor treatments. An equal number of AT2s from Sftpc-eGFP mice were seeded and cultured with either vehicle (DMSO) or under inhibitor treatments. Scale bars, 2 mm. JQ-1, a BET inhibitor. RGFP966, a Hdac3 inhibitor. A-196, a Kmt5b and Kmt5c inhibitor. **(D).** Quantification of organoid numbers following inhibitor treatment. AT2s were isolated from N = 6 or 7 mice and organoid counts were normalized to the DMSO treated control group. Error bars: mean ± SEM.

To evaluate technical screen performance, we first assessed the reproducibility of gRNA representations across experimental stages. gRNA abundance across the plasmid library, the packaged AAV library, and AT2s isolated from wild-type mice transduced with the AAV library were highly correlated (**fig. S2, B and C**, Pearson’s r > 0.97), indicating minimal bias introduced during AAV packaging and *in vivo* transduction. Moreover, AT2s harvested from different mice showed high consistency (pairwise Pearson’s correlation r > 0.92), demonstrating the robustness and reproducibility of the screening system (**fig. S2, B and C**).

To validate screen sensitivity, we examined gRNAs targeting essential genes, including *Copa*, *Prpf8*, and *Eif3a*, which showed strong depletion across multiple targeting gRNAs (**Fig. 2B**; log_2_ fold change = −1.04, −1.08, and −0.55, respectively). gRNAs targeting *Plk1*, which is essential for cytokinesis, exhibited significant depletion effects (log_2_ fold change = −0.32 and −0.28 in CM and TF screening respectively; p < 0.01, Student’s t-test; **fig. S2D**) although not reaching the current fold change cutoff for strong depletion, highlighting the stringency of our analysis. In contrast, control gRNAs, including non-targeting controls and those targeting safe harbor loci, exhibited no strong effects, confirming the specificity of the screen. Together, these results establish a robust framework for *in vivo* pooled genetic screens in the regenerating lung, which we refer to as SAGE-Peturb.

### SAGE-Perturb reveals essential epigenetic and genetic factors for alveolar repair

We observed strong deletion of gRNAs targeting histone acetylation-related genes, highlighting the potential role of this pathway in alveolar repair. Specifically, strong depletion was observed for gRNAs targeting the acetyltransferases *Kat5* (*Tip60*), *Kat8* (*Mof*), and *Taf1*, the histone deacetylase *Hdac3*, the cofactor *Ep400*, and the acetylation readers *Brd4*, *Brd2*, and *Brd8* (**Fig. 2B**). Moreover, we found gRNAs targeting *Ssrp1* and *Son* were strongly depleted, in line with their reported functions in maintaining nucleosome structure and splicing regulation of cell cycle genes, respectively^49,50^. Interestingly, the TF screen revealed marked enrichment of multiple gRNAs targeting *Nkx2-1*, a key lung lineage marker and TF essential for maintaining alveolar identity^7,43,51^.

To validate selected genes in the histone acetylation pathway, we employed the clonal AT2 organoid assay to test small molecular inhibitors targeting identified chromatin regulators. Inhibitors^52–54^ targeting Hdac3 and the bromodomain and extra-terminal (BET) proteins, including *Brd2* and *Brd4*, both of which were strongly depleted in the screen, severely impaired organoid formation, whereas inhibitors targeting non-depleted genes such as *Kmt5b* and *Kmt5c* had minimal effects (**Fig. 2, C and D**). Overall, these results demonstrate the utility of SAGE-Peturb for uncovering essential factors mediating AT2 repair in the mouse lung following IAV-mediated damage and identify histone acetylation as a previously underappreciated pathway in the regulation of alveolar repair.

### SAGE-Perturb-seq enables single cell resolution transcriptome-wide profiling of pooled genetic perturbations *in vivo*

While SAGE-Perturb successfully identified regulators of alveolar repair, its reliance on bulk analysis of limited marker gene expression, prohibiting molecular resolution into perturbation-specific cell states. To overcome this, we sought to integrate SAGE-Peturb with single-cell sequencing as a high-content phenotypic readout relating gRNA identity to transcriptomic changes at single cell resolution^32–34^.

The bulk screen highlighted *Kat8* as a previously unrecognized and essential regulator of AT2-mediated alveolar regeneration. Kat8 functions as a catalytic subunit in two distinct histone acetyltransferase complexes, the male-specific lethal (MSL) and non-specific lethal (NSL) complexes, which mediate distinct and non-redundant functions in transcriptional regulation in a wide range of physiological processes^55–59^ (**Fig. 3A**). To determine which complex underlies the regenerative requirement for Kat8, we designed a focused gRNA library targeting components of both MSL and NSL complexes and delivered it to AT2s *in vivo* via SAGE, followed by IAV infection. We harvested transduced epithelial cells (EpCAM^+^Scarlet^+^) and performed single-cell RNAseq with guide capture using 10x 5’ GEM-X chemistry methods^32,33^ (**Fig. 3B**).

**Fig. 3.**
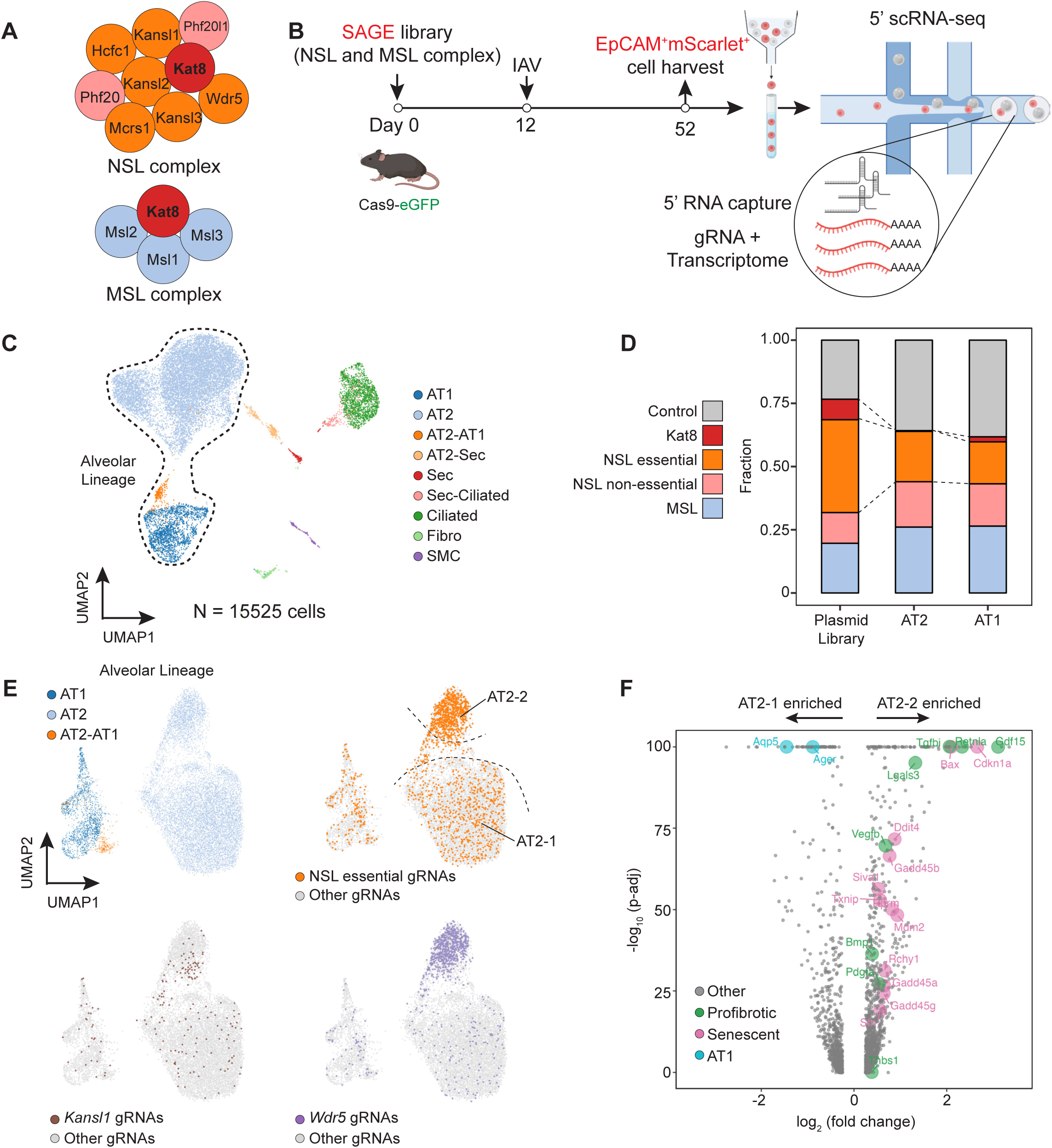
SAGE-Perturb-seq identified Kat8 as a mediator of alveolar repair through the NSL complex. **(A).** Components of the NSL and MSL complexes. Essential components of the NSL complex are colored in orange and the non-essential components in pink based on Radzisheuskaya et al., 2021. **(B).** Experimental schematic. Constitutive Cas9-eGFP mice were treated i.t. with SAGE before infection with IAV, and mScarlet^+^ epithelial cells were FACS isolated 40 days after IAV infection and subjected to 5’ scRNA-seq processing. **(C).** UMAP representation of cell populations captured in SAGE-Perturb-seq. Sec, secretory cells. Fibro, fibroblasts. SMC, smooth muscle cells. N = 15,525 cells. **(D).** Distribution of gRNA abundance across different categories of NSL and MSL complex components in the plasmid library, AT2s, and AT1s. **(E).** UMAP representation of alveolar lineage cell populations with selected cells with target gRNAs shown. **(F).** Volcano plots depicting differentially expressed genes between AT2 subpopulations. Genes with significant differences (defined as fold change > 1.5 and adjusted p-value < 0.05) are highlighted in larger circles and color-coded to indicate profibrotic, senescence-associated, and AT1 marker genes.

This generated a high quality single cell dataset, with ∼15,000 cells retained after filtering (median of 3,106 genes and 12,403 UMIs per cell, and <10% mitochondrial reads; **Fig. 3C and fig. S3A**). Our dataset recovered the major lung epithelial lineages, with a predominant representation of the alveolar lineage (AT2, AT2-AT1 and AT1, in total 82.2%; **Fig. 3C and fig. S3B**). gRNAs can be faithfully assigned to 67.4% of the cells, in line with previous reports^33^. The assignment rate varied across cell types, ranging from 10.7% in secretory cells to 73.5% in AT2s with a standard deviation of 18.3% (**fig. S3, C, D and E**). These results establish the feasibility of using SAGE-Perturb-seq for *in vivo* functional genomics to understand alveolar repair.

### Kat8 activity within the NSL complex, not the MSL complex, is essential for alveolar repair

We examined the overall changes in gRNA abundance for components of the two complexes as an indicator of their essentiality for alveolar repair. Notably, gRNAs targeting *Kat8* and essential components of the NSL complex^58^, but not those targeting components of the MSL complex, were depleted (**Fig. 3D**). To rule out the possibility that the lack of depletion was due to ineffective gRNAs targeting the MSL complex, we selected and cloned gRNAs from the library targeting distinct regions of the *Msl1* gene, an essential component of the MSL complex^60^, and assessed their genome editing efficiency. Both gRNAs demonstrated high activity, inducing insertions or deletions (indels) in approximately 80% of transduced AT2s (**fig. S4, A, B and C**). These results indicate that Kat8 and associated NSL complex, rather than the MSL complex, are critical for alveolar repair.

**Fig. 4.**
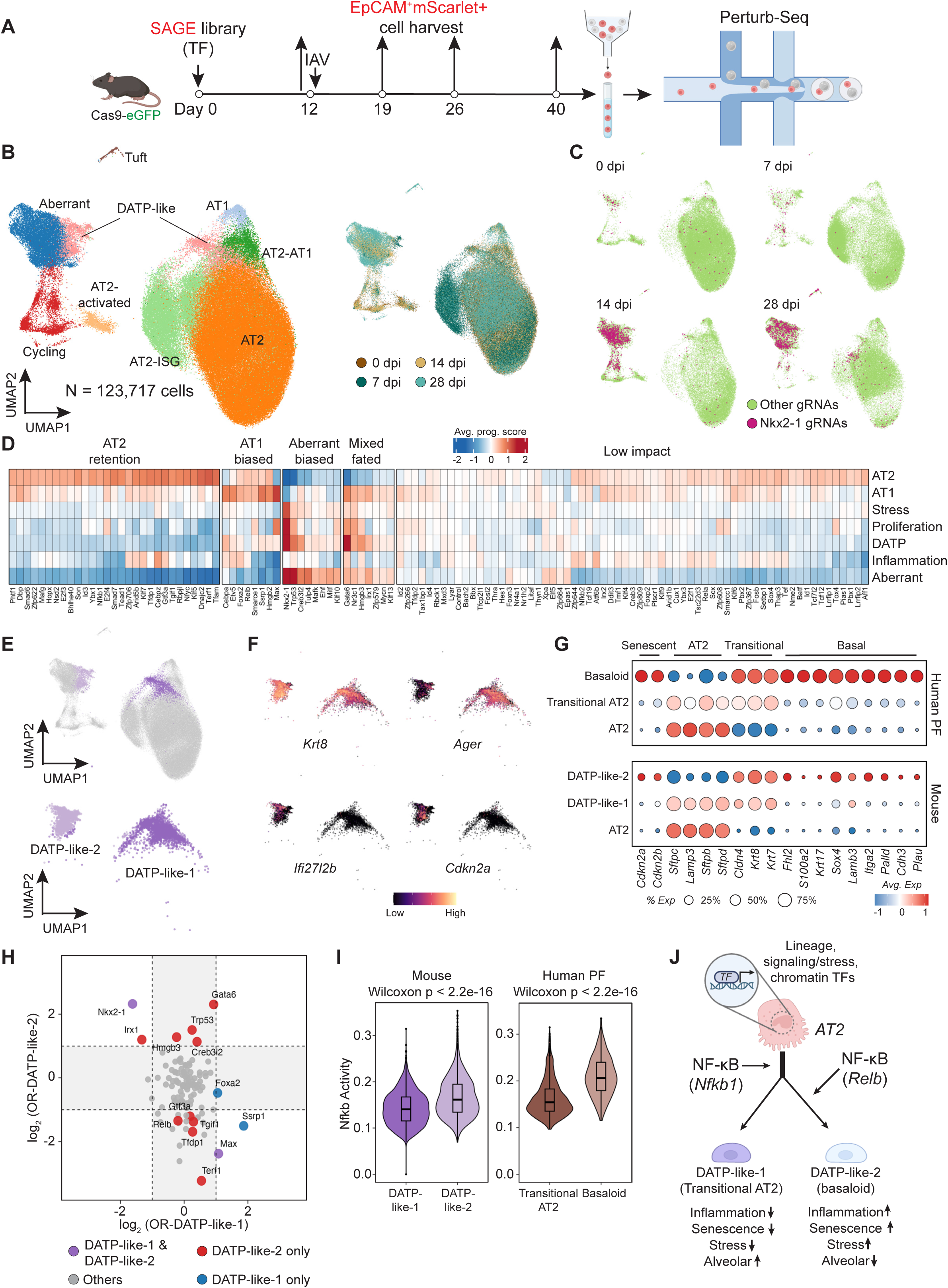
Distinct alveolar repair trajectories are governed by a network of transcription factors revealed by SAGE-Perturb-seq. **(A).** Schematic workflow of SAGE-Perturb-seq. Constitutive Cas9-eGFP mice were i.t. treated with SAGE, mScarlet^+^ epithelial cells (CD45^-^CD31^-^Pdgfra^-^EpCAM^+^) were FACS isolated before infection with IAV (0 dpi), and subsequently after infection at 7, 14, and 28 dpi. N = 2 mice per time point. **(B).** UMAP representation of the SAGE-Perturb-seq TF knockout atlas. The left panel shows the distribution of major alveolar and transitional cell populations. The right panel highlights the sample harvest time points. **(C).** Distribution of cells with *Nkx2-1* gRNAs across time points. **(D).** Heatmap of per cell program scores across gRNA target genes. TF knockout effects were grouped into AT2 retention, AT1 biased, aberrant biased, mixed fated and low impact based on dominant knockout effects. Average program scores of each TF knockouts (Methods) are centered relative to the control gRNA group and capped between −2 and 2 for visualization. **(E).** UMAP representations of the DATP-like-1 and DATP-like-2 transitional states. **(F).** Representative gene expression in DATP-like-1 and DATP-like-2 transitional states. **(G).** Dot plot visualization of gene expression similarities between mouse and human transitional states. Human PF data from Habermann et al., 2020. **(H).** gRNA abundance changes in DATP-like transitional states. Fisher’s exact test was applied to assess differential enrichment of individual gRNA knockouts relative to control gRNAs within each transitional state, with p-values adjusted using the Benjamini-Hochberg method. Significant changes, defined by log_2_ odds ratio (OR) > 1 or < −1 and FDR < 0.1 are highlighted in colored dots, indicating either shared enrichment or depletion across both DATP-like-1 and DATP-like-2 states (purple) or selective enrichment or depletion in one state (red or blue). **(I).** Violin plots of NF-*κ*B signaling activity in mouse and human transitional states. Human PF data from Habermann et al., 2020. UCell^94^ scoring was applied to compute enrichment scores for each cell. Statistical significance was assessed using the Wilcoxon rank-sum test. The center lines indicate the medians, and the boxes represent the interquartile ranges. **(J).** Schematic illustration of AT2-mediated alveolar repair regulation focusing on the exit from AT2 states and transition into distinct transitional states. Alveolar repair is jointly regulated by lineage-determining, signaling/stress-integrating, and chromatin-associated TFs. NF-*κ*B signaling influences multiple steps of AT2-mediated alveolar repair. Under conditions of inflammation, including heightened NF-*κ*B activity, the lineage trajectory is biased toward DATP-like-2 states rather than successful regeneration.

To investigate the specific role of the NSL complex in alveolar repair, we explored the transcriptional phenotypes of cells harboring NSL targeting gRNAs in the single-cell data. We first examined the distribution of gRNAs targeting NSL complex components across alveolar lineage populations. Interestingly, we discovered that a subpopulation of AT2s, AT2-2, was significantly enriched for gRNAs targeting essential components of the NSL complex compared to control gRNAs (Enrichment fold = 9.9; Fisher’s exact test p = 6.4 × 10^-203^; **Fig. 3E and fig. S4D**). AT2-2 cells also had upregulated several stress-response and senescence-associated genes, including *Cdkn1a*, *Gadd45a/b/g*, *Sfn*, and *Ddit4*, indicative of the activation of an early senescence-like program (**Fig. 3F**). Moreover, these cells also showed downregulation of AT1 markers such as *Aqp5* and *Ager*. These findings suggest that loss of NSL function triggers a senescence-like state and blocks progression towards AT1 fate.

Furthermore, AT2-2 cells exhibited increased expression of profibrotic growth factors such as *Gdf15* and *Pdgfa*^61,62^, as well as secreted factors associated with tissue remodeling like *Retnla*^63^, accompanied by reduced expression of anti-fibrotic genes including *Apoe* and *Lcn2*^64,65^ (**Fig. 3F and fig. S4E**). This suggests that a loss of NSL complex function in AT2s not only compromises their proliferative and differentiative capacity but may also perturb epithelial-fibroblast communication following injury through a pro-fibrotic secretome^66–68^.

### SAGE-Perturb-seq uncovers transcription factor networks driving alveolar repair

With SAGE-Perturb-seq established, we next sought to systematically compare TF function in alveolar repair. We assembled a 260 gRNAs TF library, including gRNAs targeting 118 TFs (2 gRNAs each), alongside control gRNAs targeting safe harbor loci and intergenic regions (**table S1**). To capture the full dynamics of the cell types and states, we performed SAGE-Peturb-seq at four key time points during repair: directly prior to IAV infection (0 dpi), peak infection (7 dpi), peak proliferation (14 dpi), and following resolution (28 dpi) (**Fig. 4A**). Following quality control filtering, exclusion of non-alveolar lineages (e.g., ciliated cells), and implementation of a stringent dominant gRNA assignment strategy (**Methods**), we retained approximately 123,000 high-quality single-cell perturbation profiles spanning the full time course with single gRNA assignment rate of 46.9% (123,717/263,587) across all time points (**Fig. 4B and fig. S5A**). Perturbation profiles were recovered for all 118 targeted TFs, with 116 TFs represented by more than 100 confidently assigned cells (**fig. S5B**).

Our single cell transcriptome libraries captured dynamic shifts in epithelial cell states across time points, consistent with the phases of injury and repair following IAV infection^39^ (**Fig. 4B and fig. S5, C and D**). AT2 recovery was lowest at 7 dpi and gradually restored by 14 and 28 dpi, mirroring the expected trajectory of IAV-induced epithelial damage and subsequent repair. A transient interferon-responsive AT2 population (AT2-ISG), characterized by elevated expression of interferon-stimulated genes, emerged at 7 dpi and declined over time, reflecting the peak and resolution of inflammation. In contrast, AT1s were underrepresented across time points compared to AT2 cells^39,69^, likely due to low recovery during single-cell processing.

We identified an aberrant cell population marked by expression of *Tff2, Olfm4, Gkn3,* and *Agr2*, consistent with a secretory-like, metaplastic phenotype. In line with prior studies^7,70^, this aberrant *Tff2*+ secretory-like population was significantly enriched for cells harboring *Nkx2-1* gRNAs compared to control gRNAs (Fisher’s exact test, odds ratio = 15.9, false discovery rate (FDR) < 2.2 × 10^-16^; **Fig. 4C**). In parallel, *Nkx2-1* knockout cells became the most expanded epithelial population by 28 dpi (**fig. S5E**) and retained MHCII expression (**fig. S5C**), indicating that loss of *Nkx2-1* not only biases cells toward an aberrant fate but also promotes their expansion in the context of injury.

### Functional mapping of transcription factor knockout effects during alveolar repair

Capturing the full spectrum of proliferative, transitional, and aberrant cell states is essential for mapping TF knockout effects during alveolar repair. However, traditional approaches relying on discrete cell type clustering may obscure the underlying transcriptional heterogeneity and fail to resolve the dynamic trajectories embedded in the data. To overcome this limitation, we applied HOTSPOT^71^, a cell type-agnostic algorithm that identifies locally co-varying gene modules, to reveal subtle and continuous transcriptional programs.

Applying HOTSPOT analysis, we identified 25 gene programs, which we consolidated into 14 meta-programs based on their functional similarity and enrichment across specific cell states (**fig. S6A**). These fell into two broad categories: cell type/state programs, which define stable or transitional identities, and cell activity programs, which reflect dynamic cellular responses layered on top of identity. Among the cell type/state programs, we observed signatures for AT2, AT1, DATP, tuft, and aberrant cells. The cell activity programs included programs related to cell proliferation, inflammatory and stress responses, cytokine secretion and checkpoint regulation, cytoskeletal remodeling, barrier-signal integration, redox-lipid damage control, and housekeeping.

To assess the functional relevance of these programs, we next asked whether differences in their average scores (**Methods**) could capture known TF knockout effects. Using the AT1 program as a test case, we calculated the average program score across cells harboring gRNAs targeting individual TFs. *Nkx2-1* knockout cells exhibited reduced AT1 program scores (**fig. S6B**), indicating that *Nkx2-1* is critical for promoting AT1 differentiation, in line with previous studies^51^. In contrast, knockouts of *Max*, *Ssrp1*, *Cebpa*, *Hmgb2*, *Etv5*, and *Gata6* led to elevated average AT1 program scores compared to other TF knockouts. At the gene level, cells lacking *Etv5*, *Cebpa*, *Gata6*, *Ssrp1*, or *Max* exhibited upregulation of canonical AT1 markers, including *Ager*, *Hopx*, *Vegfa*, *Aqp5*, *Cav1* and *Spock2*^43,72^ (**fig. S6C**). At the cellular level, these knockouts also shifted a fraction of cells toward the AT1s (**fig. S6D**). Supporting this, prior studies have reported enhanced AT1 differentiation upon deletion of *Cebpa* or *Etv5* in mouse lung injury models^8,9^. Together, these results validate transcriptional program analysis using average program scores as a scalable framework for interpreting TF knockout effects, recovering known regulators of alveolar fate such as *Nkx2-1*, *Cebpa*, and *Etv5*, and uncovering additional candidates, such as *Gata6*, *Max*, *Ssrp1* and *Hmgb2*, that may restrain AT1 differentiation during lung repair.

### Defining functional groups of transcription factors by knockout effects in alveolar repair

To provide a systems-level view of the transcriptional regulatory network underlying alveolar repair, we next grouped TF functions based on their knockout effects across gene program activity profiles. We focused on seven programs central to alveolar repair, including cell state programs (AT2, AT1, DATP, and aberrant states) and activity programs related to proliferation, inflammatory and stress response (**fig. S7A**). To identify shared patterns, we first applied unsupervised K-means clustering to group TFs based on their multi-program response profiles (**fig. S7, B and C**). Building on the effects of these clusters, we applied a supervised rule (**Method**) to assign each TF knockout to a dominant effect category based on average program score shifts in comparison to control gRNAs toward AT2, AT1, or aberrant states. Knockouts that strongly upregulated the AT2 program were classified as AT2 retention, while the remainder were assigned as (1) AT1-biased, if knockouts elevate AT1 program scores without activating or minimally affecting aberrant or stress pathways; (2) aberrant-biased, if knockouts strongly upregulate aberrant and stress signatures; (3) mixed fated, if knockouts upregulated both AT1 and aberrant programs; or (4) low impact (**Fig. 4D**). This approach enables systematic interpretation of TF perturbations in terms of their influence on epithelial cell-state dynamics, providing a conceptual scaffold for understanding how distinct TFs may cooperatively shape, bias, or destabilize epithelial repair trajectories following injury.

We hypothesized that TFs exhibiting similar knockout effects may share co-regulatory relationships. To test this, we focused on TFs whose knockout led to upregulation of AT1 programs, including *Foxa2, Cebpa, Etv5,* and *Max*. Loss of these factors upregulated AT1 programs, suggesting that they may contribute to maintaining AT2 identity and restraining premature AT1 differentiation. Based on this observation, we next asked whether these TFs could physically co-bind regulatory regions of AT1 program genes as a proxy for co-regulation. Using publicly available ChIP-seq data and ATAC-seq profiles from mouse AT2s, we analyzed open promoter regions (−2 kb to +0.5 kb around transcription start sites, TSS) of AT1 genes and found significant co-binding relative to background (**fig. S7D**). Complementing this, HOMER motif analysis of ATAC-seq-defined promoters and putative enhancers revealed significant enrichment of Cebpa and Foxa family motifs near AT1 program genes (**fig. S7E**). These findings support the functional coherence of this group of TFs and highlight the power of SAGE-Perturb-seq to uncover biologically meaningful co-regulatory networks in alveolar repair.

### SAGE-Perturb-seq identifies reparative and pathological transitional cell states in alveolar repair

Transcriptional program analysis revealed that the DATP-like population was not homogenous, but instead composed of two transcriptionally distinct states (**Fig. 4E and fig. S8A**). These states, termed DATP-like-1 and DATP-like-2 here, share gene expression signatures with the previously described DATPs, also referred to as ADIs or PATS^6,14,15^. Both states expressed canonical DATP markers, such as *Krt8* and *Cldn4* (**Fig. 4, E, F and G**), however, they diverged in their broader gene expression profiles (**fig. S8B**). DATP-like-1 cells retained expression of core alveolar lineage markers including *Sftpc*, *Abca3*, *Aqp5*, and *Ager*, consistent with a trajectory toward differentiation into mature alveolar epithelium. In contrast, DATP-like-2 cells exhibited enrichment for genes involved in interferon and innate immune signaling (*Ifit1/2/3*, *Stat1*, *Irf1/7*), endoplasmic reticulum and oxidative stress responses (*Gsto1*, *Gstm1/2/5*), as well as signatures associated with cellular senescence (*Cdkn2a*, *Cdkn2b*) and apoptosis (*Bok*, *Pmaip1*), suggestive of an inflammation- and stress-induced state.

Comparison with single-cell profiles from diseased human lungs revealed that these two states closely parallel transitional epithelial populations observed in human pulmonary fibrosis samples. DATP-like-1 cells resemble the transitional AT2 population (also known as alveolar basal intermediate 1, ABI), which has been proposed as a reparative intermediate. DATP-like-2 cells mirrored the basaloid population (also known as ABI2 or KRT17^+^/KRT5^-^ cells), a pathological state that expands in fibrotic lungs such as those of pulmonary fibrosis (PF) patients^17–19,21^ (**Fig. 4G and fig. S8, C and D**). Our data demonstrate that both reparative and aberrant transitional states can be recapitulated in a genetically-perturbed, injury-responsive, physiologically-relevant mouse model, establishing a novel experimental framework for investigating the regulation of these transitional states.

To determine whether DATP-like-1 and DATP-like-2 states arise sequentially or represent distinct transitional routes of AT2 in response to injury, we performed trajectory inference analyses using Slingshot^73^. This revealed that DATP-like-1 was strongly biased toward AT1 differentiation, whereas DATP-like-2 was biased toward the aberrant branch (**fig. S8E**), indicating that they are parallel, rather than sequential states. Together, our findings suggest that AT2s can independently enter either a repair-biased DATP-like-1 state or a maladaptive, stress-primed DATP-like-2 state, the latter of which may give rise to an aberrant population.

### SAGE-Perturb-seq identifies genetic players that shape different transitional states

The DATP-like-1 and DATP-like-2 states represent murine counterparts of the human transitional AT2 and basaloid states^17–19,21^, respectively. These populations have emerged as critical nodes in the balance between effective regeneration and pathological remodeling in lung diseases. Despite their clinical relevance, the regulatory mechanisms governing their emergence remain unclear. To define the genetic logic underlying these trajectories, we used SAGE-Perturb-seq to map how targeted perturbations bias AT2 fate toward reparative or pathological transitional states.

We quantified gRNA enrichment and depletion relative to control gRNAs within DATP-like-1 and DATP-like-2 states, thereby nominating candidate regulators that bias AT2 fate toward each trajectory. This analysis identified lineage and differentiation TFs (*Nkx2-1*, *Foxa2*, *Gata6*, *Irx1*), signaling and stress response TFs (*Tgif1*, *Relb*, *Trp53*, *Creb3l2*), as well as chromatin and transcriptional machinery (*Gtf3a*, *Ssrp1*, *Hmgb3*, *Terf1*) that collectively shape the emergence of these two states (**Fig. 4H**).

Among these, Nkx2-1 emerged as a pivotal bifunctional regulator, with gRNAs targeting *Nkx2-1* significantly depleted in DATP-like-1 but enriched in DATP-like-2. This inverse pattern aligns with the well-established role of Nkx2-1 in driving functional alveolar repair and facilitating AT2 to AT1 differentiation^51^. Its loss impairs this trajectory, leading to reduced entry into the reparative DATP-like-1 state and a corresponding accumulation of cells in the DATP-like-2.

In contrast, *Max* knockout had reciprocal effects, with gRNAs enriched in DATP-like-1 but depleted in DATP-like-2. As Max is the obligate binding partner of Myc and required for Myc-driven transcriptional programs^74^, its loss is expected to blunt Myc activity. Indeed, DATP-like-1 cells displayed significantly lower Myc program activity compared with DATP-like-2 cells (Wilcoxon p < 2 × 10^-16^; **fig. S8F**), suggesting that higher Myc activity is required for DATP-like-2 emergence. Consistent with this, knockout of *Tfdp1*, a direct Myc target^75^, also exhibited reduced representation in the DATP-like-2 state (**Fig. 4H**). Together, these results indicate that DATP-like-2 emergence depends on the Myc/Max axis, and that suppression of this pathway diverts AT2s toward the DATP-like-1 trajectory.

NF-*κ*B-related TFs govern the emergence of the DATP-like-2 state. *Relb* knockout markedly depleted this population, indicating a requirement for non-canonical NF-*κ*B signaling. Consistent with this, *Relb*-deficient cells showed a bias toward AT1 fate (**Fig. 4D**), favoring a reparative rather than the aberrant trajectory. We also observed that *Rela* perturbation produced a similar, though weaker, trend toward DATP-like-2 depletion (Fisher’s exact test, odds ratio = 0.42; FDR = 0.34), suggesting that canonical NF-*κ*B activity may likewise contributes to DATP-like-2 emergence. Supporting this, canonical NF-*κ*B activity, inferred from its gene expression signature, was elevated in DATP-like-2 cells and in the human basaloid population compared with their reparative counterparts (**Fig. 4I**). Interestingly, canonical NF-*κ*B signaling also influences AT2 progenitor exit, as *Nfkb1* knockout resulted in retention of the AT2 signature and impaired entry into the regenerative trajectory (**Fig. 4D**). Beyond NF-*κ*B signaling, other inflammatory regulators were similarly engaged. IFN-gamma/STAT1 and STAT3 pathways were also elevated in DATP-like-2 and human basaloid populations (**fig. S8G**). These findings indicate that elevated NF-*κ*B (through both canonical and non-canonical arms), along with IFN-gamma/STAT1 and STAT3 activity, may represent a shared regulatory node that drives the emergence of pathological transitional states in both acute and chronic lung injury.

Taken together, these findings show that the emergence of distinct DATP-like states is orchestrated by diverse genetic regulators acting within inflammatory niches. These results also demonstrate the power of SAGE-Perturb-seq to resolve cell-state-specific mechanisms *in vivo*. This integrative framework provides a lens to dissect how regenerative versus pathological epithelial trajectories are differentially specified during tissue repair.

## Discussion

Effective lung repair following viral respiratory infection is critical for host survival. Incomplete or aberrant repair can lead to a variety of post-viral sequelae, many of which are driven by a weakened or dysfunctional epithelial barrier after viral clearance^40,76,77^. Understanding the molecular drivers of alveolar repair is therefore key to identifying new therapeutic opportunities.

Here, we present the first *in vivo* genetic screening platform for the lung epithelium in its native context, enabling a high-resolution dissection of transcriptional and epigenetic regulation of alveolar repair after viral injury. SAGE-Perturb allowed us to identify modulators of histone acetylation as essential components for AT2 proliferation. However, this screening mode uses cell surface markers to define cell populations, thus obscuring potential cellular heterogeneity. To overcome this limitation, we coupled it with single-cell profiling in SAGE-Perturb-seq, which uncovered the Kat8-associated NSL complex as a key regulator of repair. Its loss triggered AT2 senescence-like phenotypes and induced a profibrotic secretory program. It is tempting to speculate that this process may promote local fibrosis and thus exacerbates IAV pathology. Intriguingly, genetic variants of *KANSL1*, a core NSL complex component, have been associated with idiopathic pulmonary fibrosis (IPF)^78–80^, a condition also characterized by impaired alveolar regeneration. These findings suggest that the NSL complex is critically involved in AT2-driven alveolar repair and thus may represent a target with potential therapeutic implications in pulmonary fibrosis and other lung pathologies with impaired alveolar regeneration.

Recent studies have described an AT2 to AT1 transitional state, termed as DATP, ADI, or PATS^6,14,15^. However, its relationship to the distinct intermediates observed in human pulmonary fibrosis, the reparative transitional AT2 state and the pathological basaloid state^17–19,21^, has remained unresolved, raising the question of whether the basaloid signature can be reproduced in mouse models. A recent study has shown that basaloid-like states can emerge through *Sftpc* mutant expression in a mouse model^81^. Here, using an IAV-induced lung injury model, we demonstrate that two distinct transitional states observed in human lungs can also be recapitulated as a reparative DATP-like-1 population and a pathological DATP-like-2 population. This observation was likely enabled by the large number of epithelial cells we profiled and the amplification of the DATP-like-2 state under genetic perturbations. These findings provide additional rationale for the use of IAV to model aspects of early lung fibrosis. The ability to distinguish these two states suggests therapeutic potential to selectively target the maladaptive state without compromising the regenerative state.

Leveraging SAGE-Perturb-seq, we were able to further dissect the regulatory networks driving DATP-like-2 emergence. We found that elevated inflammatory signaling, such as through NF-*κ*B (via *Relb* and *Rela*), diverts AT2s into the pathological transitional state. In addition, TF knockouts such as *Nkx2-1*, *Gata6*, *Trp53*, *Irx1*, *Hmgb3*, and *Creb3l2* accumulated preferentially within DATP-like-2 cells. This supports a model in which inflammatory cues promote AT2 entry into the DATP-like-2 state, while genetic perturbations (or somatic mutations) may help stabilize this state or restrict its reversal. Given the similarities between the DATP-like-2 and basaloid states and the accumulation of these cells in pathologic conditions, it is likely that inflammation and genetic vulnerability accelerates the persistence and expansion of basaloid populations, thereby amplifying fibrotic remodeling and disease progression. This is consistent with recent observations of macrophage accumulation in fibrotic niches as a source of sustained inflammatory signaling^40,82^. Our insights highlight the need to re-evaluate early anti-inflammatory interventions in pulmonary fibrosis onset or exacerbation.

SAGE enables interrogation of gene function using flexible phenotypic readouts, ranging from simple bulk proliferation to high-resolution single-cell whole-transcriptome profiling. It supports screens of 100-200 genes using 4-5 mice, and is easily scalable to ∼1,000 genes, accommodating libraries such as the Kinome^83^ and Phosphatome^84^. With advances in CRISPR technology, including the development of Prime Editor mouse models ^85^ and the adaptive tropism of AAVs, this platform can be easily repurposed to assess variant-specific contributions to disease phenotypes and enable functional genomic studies in other tissues. Together, these approaches and findings create a foundation for future gene function studies in the mouse lung and open avenues for rationally targeting transitional states to improve recovery after acute or chronic injury.

## Acknowledgments

We thank Jayaraj Rajagopal, Siddhartha Jena, Jason Buenrostro, Carla Kim, Jiaqi Zhang, Sebastiano Cultrera di Montesano, and all members of the Chen and Franklin Labs for helpful discussions. We thank Yifan Zhang, Ruth Raichur and Somkene (DK) Alakwe for help in experiments. We thank David Ziehr, Forest (Yunhan) Xu and Raghu Chivukula for kindly sharing the Sftpc-eGFP mice.

D.S. acknowledges support from the NHLBI Pathway to Independence Award (K99HL181185) and the American Lung Association Catalyst Award (CA-1447794). This work was supported by the National Institutes of Health (grant nos R01HG010647 to F.C.). F.C. acknowledges support from the Searle Scholars Award, the Burroughs Wellcome Fund CASI award, the Merkin Institute, Harvard Stem Cell Institute, and the NYSCF. F.C. is an NYSCF Roberston Investigator. R.A.F. acknowledges support from the NIH Maximizing Investigators’ Research Award (R35, GM150816), the Charles H. Hood Foundation, and the Harvard Stem Cell Institute.

## Author contributions

D.S., D.A.H., R.A.F., and F.C. conceived of the project. D.S., D.A.H, D.S., B.O., S.B., and G.M. performed experiments. D.S., D.A.H, C.H., and J.A.W. performed analyses. D.S., D.A.H., R.A.F., and F.C. interpreted the results. D.S., D.A.H., R.A.F., and F.C. wrote the manuscript with input from all authors.

## Data and code availability

The raw data and processed data generated in this study will be submitted to NCBI Gene Expression Omnibus. The analysis pipeline will be deposited in the GitHub repository. Processed data will be deposited to the Single Cell Portal. Plasmids generated in this study will be deposited to Addgene.

## Materials and methods

### Mice

All animal experiments were performed in accordance with institutional regulations after protocol review and approval by the Institutional Animal Care and Use Committee (IACUC) at the Broad Institute (protocol 0211-06-18-2) or Harvard Medical School (protocol IS00003152). Mice were bred and/or housed in a specific pathogen-free facility. C57BL/6J (RRID:IMSR_JAX:000664), constitutive Cas9-eGFP (RRID:IMSR_JAX:026179), Spc-CreER (RRID:IMSR_JAX:028054), LSL-Cas9-eGFP (RRID:IMSR_JAX:026175) and Ai14 (RRID:IMSR_JAX:007914) mice were obtained from Jackson Laboratories. Female mice aged 8-12 weeks were used for all experiments.

### Tamoxifen administration

Tamoxifen (Merck, T5648) was dissolved in corn oil (Merck, C8267) to prepare a 20 mg/mL stock solution. Mice received tamoxifen via intraperitoneal (IP) injection at a dose of 0.2 mg/g body weight. The number of injections and timing of administration are specified in the corresponding experimental scheme figures.

### Immunostaining

Mouse lung tissues were collected and fixed overnight at 4°C in 4% paraformaldehyde (PFA). Fixed tissues were cryoprotected, embedded, and sectioned at 12–20 μm thickness for immunofluorescence analysis. Lung sections were rinsed once in PBS and permeabilized with 0.3% Triton X-100 in PBS for 10 minutes at room temperature (RT). After three PBS washes, sections were blocked in 0.5% (w/v) bovine serum albumin (BSA) and 0.1% Triton X-100 in PBS (blocking buffer) for 1 hour at RT.

Primary antibodies, proSPC (1: 500, Merck, Ab3786), were diluted in blocking buffer and incubated overnight at 4°C. After washing, secondary antibodies or conjugated primary antibodies diluted in blocking buffer were applied overnight at 4°C. The following secondary antibodies were used: donkey-anti-rabbit Alexa 594 (1: 1000, Thermo Fisher Scientific, A32754) and donkey-anti-rabbit Alexa 647 (1: 1000, Thermo Fisher Scientific, A31573). The following conjugated primary antibodies were used: eGFP antibody (1: 500, SySy, N0304-At488-L) and RFP antibody (1: 500, SySy, N0404-AF568). The next day, nuclei were stained with DAPI (Sigma, D9542) in PBS for 10 minutes at RT. After three PBS washes, sections were mounted using Fluoromount™ Aqueous Mounting Medium (Merck, F4680).

Images were acquired using a Nikon Ti2 inverted research microscope with spinning disk confocal (Andor Dragonfly), collecting z-stacks using a 40× objective. Representative single-plane images are shown. Image processing was performed using ImageJ (version 2.3.0).

### Organoid culture

#### For small molecule inhibitors

Mouse lungs were cleared by perfusion with cold PBS through the right ventricle, and minced into small pieces. Tissues were dissociated with 4 ml of dissociation mixture containing Collagenase I (1mg/ml, Merck, C9891), Dispase II (2U/ml, Merck, 4942078001) and Dnase I (0.1 mg/ml, Merck, 11284932001) at 37 °C for 35 min with horizontal shaking at 140 rpm (after 20 min of incubation, the mixture was triturated with P1000 pipette to facilitate dissociation). The cell suspensions were then filtered through 40-μm cell strainers, diluted with 8 ml of wash medium (Advanced DMEM/F12 (Thermo Fisher Scientific, 12634010), supplemented with 10mM HEPES (Thermo Fisher Scientific, 15630080), 1x GlutaMAX (Thermo Fisher Scientific, 35050061), 50 U/ml Pen/Strep (Thermo Fisher Scientific, 15070063) and 10% FBS (Merck, F4135)) and centrifuged at 600*g* for 8 min at 4°C. The cell pellet was resuspended in 2ml of RBC lysis buffer (Thermo Fisher Scientific, 00-4333-57) and lysed for 2 min at room temperature. 10 ml of wash medium was added to terminate the reaction, followed by centrifugation at 600*g* for 8 min at 4°C. The cell pellet was resuspended in a staining buffer (PBS with 0.5%BSA and 2mM EDTA) for further staining (when testing SAGE integration) or directly proceeding to sorting (when using Sftpc-eGFP mice or Sftpc-CreER;LSL-tdTom mice). The antibodies used were as follows: CD45 (30-F11)-APC (BD Biosciences), CD31 (MEC13.3)-APC (BD Biosciences), EpCAM (G8.8)-PE-Cy7 (BioLegend) and MHC-II (I-A/I-E, M5)-FITC (eBiosceince). Sony MA900 cell sorter was used for the sorting at Broad Institute Flow Cytometry Core.

#### For quantification of integration efficiency

Mice were sacrificed using a lethal dose of ketamine/xylazine and digested in a dispase/liberase solution as previously described^86^. Briefly, lungs were perfused with 5mL of 5mM EDTA in PBS, inflated with dispase (Corning), before being chopped and digested with a liberase (Sigma-Aldrich) and DNAse (Sigma-Aldrich) mixture in RPMI at 37°C on an orbital shaker. Lung digests were neutralized with FBS (GeminiBio) and filtered into a single cell suspension through a 70um filter before proceeding to staining. AT2s were gated as live singlets CD45^-^CD31^-^Pdgfra^-^EpCAM^+^CD104^-^MHCII^+^, and sorted as either Scarlet^-^ or Scarlet^+^.

AT2 organoids are cultured as previously described^87,88^. Briefly, 10,000 cells are seeded in 50 μl Matrigel domes in a 24-well plate well. Each well is cultured with 600 μl Advanced DMEM/F12 as base medium, supplemented with 10mM HEPES, 1x GlutaMAX, 50U/ml Pen/Strep, N-Acetylcysteine (Merck, A9165, 1.25mM), 1× B27 (ThermoFisher, 12587010), 1× ITS (ThermoFisher, 41400045), 1× Antibiotic-Antimycotic (ThermoFisher, 15240062), hFGF7 (ThermoFisher, 100-19, 50ng/ml), mNoggin (ThermoFisher, 250-38, 25ng/ml), hFGF10 (ThermoFisher, 100-26, 25ng/ml), mEGF (ThermoFisher, 315-09, 50ng/ml), SB431542 (Biotechne, 1614, 10μM), BIRB796 (Biotechne, 5989, 1μM), CHIR99021 (Biotechne, 4432, 3μM). For the first 48hrs of culture, Y-27632 (Biotechne, 1254, 10μM) is supplemented. Organoids are cultured for 12 days when colony numbers are counted.

### Cloning

AAV vector is linearized by enzymatic digestion using BsaI-HFv2 (New England BioLabs, R3733L) at 37 °C overnight, followed by gel purification with Macherey-Nagel Gel and PCR clean-up kit (Macherey-Nagel, 740609).

For the chromatin modulator screen, 3 gRNA sequences targeting each gene were randomly selected from (Shi et al., 2015). The target gene and gRNA sequences are listed in table S1.

For the MSL and NSL complex focused screen, components of the two protein complexes were curated based on Radzisheuskaya et al., 2021. gRNA sequences were designed using Broad Institute online gRNA designing tool (https://portals.broadinstitute.org/gppx/crispick/public). For each component, five gRNAs were selected. In addition, 10 non-targeting control gRNAs and 10 gRNAs targeting intergenic regions were included as controls. gRNA sequence details are in table S1.

For the transcription factor screen, gene expression enrichment in AT2 cells, AT1 cells and different transitional cell populations were computed using seurat FindAllMarkers function from publicly available single cell RNA-Seq datasets^6,47,89^. Transcription factors and their enrichment are listed in table S1. gRNA sequences were designed using Broad Institute online gRNA designing tool (https://portals.broadinstitute.org/gppx/crispick/public). For each transcription factor, five gRNAs were selected. In addition, 30 non-targeting control gRNAs and 30 gRNAs targeting intergenic regions were included as controls. gRNA sequence details are in table S1.

Twenty-five nucleotide homology sequences flanking the BasI cleavage sites of the AAV plasmid were added at 5′ and 3′ ends of the gRNA sequences for cloning. Oligos were purchased as oPools™ Oligo Pools from IDT at 1 pmol per oligo scale. 100 ng of linearized vector was mixed with the oligo pool in a molar ratio of 1: 200 and with NEBuilder HiFi DNA Assembly Master Mix (New England BioLabs, E2621) according to the manufacturer’s instructions. The mixture was incubated at 50°C for 1 h and cooled down on ice. 8-10 transformation reactions were performed with 2 μl of the reaction mixture in 50 μl Stellar™ Competent Cells (636763, Takara). A small fraction (20 μl) of the competent cells was plated to calculate the amount needed for 500X representation per gRNA as seeding in Maxiprep. At the same time, the same amount of linearized vector was transformed in the same way for background control (colony number < 5% of the library construction transformation). gRNA library pool plasmid was harvested using ZymoPURE II Plasmid Maxiprep Kit (Zymo, D4203). gRNA library quality was subsequently checked by next-generation sequencing (NGS). Primer sequences used are listed in table S1.

### AAV packaging and purification

AAVs were produced in HEK293FT cells (Thermo Fisher Scientific, R70007, within 10-15 passages) and purified by iodixanol gradient centrifugation. In brief, HEK293T cells were expanded in Advanced DMEM/F12 (Thermo Fisher Scientific, 12634010), supplemented with 10mM HEPES (Thermo Fisher Scientific, 15630080), 1× GlutaMAX (Thermo Fisher Scientific, 35050061), 50 U/ml Pen/Strep (Thermo Fisher Scientific, 15070063) and 10% FBS (Merck, F4135). 24 h before transfection, cells were seeded in 150 mm dishes (Thermo Fisher Scientific) at 12 million cells per dish culture with 25 ml medium. Cells were transfected with 12 μg of pAdDeltaF6 plasmid (Addgene #112867), 10 μg of AAV2/9 serotype plasmid (Addgene #112865) and 6 μg of AAV plasmids using Transporter 5 transfection reagents (Polysciences, 26008). 72-96 h after transfection, HEK293FT cells were detached with cell scraper and collected in 250mL Centrifuge Tubes. Cells were centrifuged at 1500*g* for 15 min. Then, cells were resuspended in 1.5-2 ml of 1× PBS solution per 5 dishes of cell pellets and incubated on ice. 20 μl of protease inhibitor PMSF (Merck, 93482) was added to cell suspension and cells were lysed with a sonicator with microtips at 30 amplitude for 2x 10s. Another 20 μl of protease inhibitor PMSF was added. Then, 50 μl of 100mM MgCl2 was added together with 50 μl of 10mg/ml DNAse I (Merck, 11284932001) to digest genomic DNA at 37 °C for at least 30 min. Iodixanol gradients were prepared by sequential pipetting of the following iodixanol solutions: 1.5 ml (17%), 1.5 ml (25%), 2 ml (40%) and 2 ml (60%) into a 8.9 ml OptiSeal Polypropylene Tubes (Beckman Coulter, 361623). Cell lysates were carefully loaded on top of the 17% iodixanol layer. Gradients were ultracentrifuged using the Beckman type 70.1 Ti rotor at 350,000 rcf for 2.25 h at 18 °C. To recover the AAV particles, 500 μl fractions were taken from the tube and stored in deep 96 well plates. Viral genomes were quantified using qPCR (details below) and the 4 fractions with the highest viral load were collected, diluted with and with PBS and centrifuged through a 15 ml Amicon 100 kDa MWCO filter unit (Amicon) at 4,000*g* for 30 min. These dilution and centrifugation steps were repeated for three rounds. The resulting AAV solutions were quantified again using qPCR (details below) and aliquoted to store at 4 °C for short-term use (< 1 week) or store at −80 °C for long term. The AAV particle concentration was determined by qPCR. In brief, 5 μl of the fraction after ultracentrifugation, isolated AAVs or Addgene AAV standards (Adgene #37825-AAV9) were treated with with DNase I (Merck, 11284932001) at 37 °C for at least 1 hr, followed by Proteinase K digestion at 56 °C for at least 2 hrs before preparing a threefold serial dilutions with ddH2O. qPCR reactions with primers targeting the WPRE region (WPRE_F, CAATCCAGCGGACCTTCCTT; WPRE_rev, AGATCCGACTCGTCTGAGGG) for normal AAVs and the mScarlet region (mScarlet_F, AAGAAGCCCGTGCAGATGCC; mScarlet_R, GCCACGGAGCGTTCGTACTG) for SAGE were performed with 3 μl of the diluted AAV template, 5 µl Power SYBR™ Green PCR Master Mix (Thermo Fisher Scientific, 4368706), 2.5 µM of each primers in a total of 2 µl. qPCR was performed using RioRad CFX 384 system and at conditions as follows: (1) 95 °C for 10 min; (2) 40 cycles of 95 °C for 15 s and 60 °C for 1 min (40 cycles).

### Influenza A Virus

A/Puerto Rico/8/1934 (H1N1) (Charles River Laboratories) was used for all influenza A virus (IAV) infections. Infectious dose was determined by tissue culture infectious dose 50 (TCID_50_), as described previously^90^. Briefly, MDCK cells were infected with viral dilutions for 48 hours at 37 °C. TCID_50_ was determined by hemagglutination assay with chicken red blood cells (Rockland) and calculated using the Reed-Muench method^91^.

### In vivo pooled genetic screen

To estimate the screenable library size *in vivo*, we considered that previous studies have estimated that ∼1-2 million AT2 cells can be isolated from an adult mouse lung^86^. With a transduction efficiency of ∼20% at the dose of 1 × 10^11^ vg, we estimate that approximately 200,000-400,000 recoverable AT2 cells are transduced *in vivo*. Assuming a minimum of 500 cells per gRNA is required to ensure robust screening performance, this setting could support a library of up to 400-800 gRNAs, corresponding to approximately 100-200 genes (3-5 gRNAs per gene).

AAV particles carrying gRNAs library for chromatin modulators or transcription factors genes were generated as described in the ‘AAV production and purification’ section. A single dose of 1 × 10^11^ viral particles in a total volume of 50 μl was administered through intratracheal delivery (i.t.). Two weeks after AAV gRNA delivery, mice were infected with 150 TCID_50_ of IAV. Mice were anesthetized with a ketamine/xylazine mixture and 30ul of viral inoculate was instilled intranasally. Mice were monitored for weight loss before lung tissue harvesting and processing.

### Flow cytometry and fluorescence activated cell sorting

Mouse lungs were harvested and digested as described in organoid assay for quantification of integration efficiency. For flow cytometry, approximately 1 million cells fromwhole lung suspension were stained/well of a 96-well plate. Cells were stained with Fc block (1:250) and live/dead stain (1:150) for 10 minutes at room temperature in the dark before addition of antibody cocktail and incubated for 30 minutes on ice in the dark. Cells were washed three times with 1% FBS in PBS before data collection. Flow cytometry samples were collected on a Symphony A5 or A3 (BD). For cell sorting, the entire lung homogenate was stained using 1 million cells/well in a 96-well plate. Fc block and antibody cocktails were stained for 30 minutes on ice before washing. Before sorting, 1:5000 DAPI stain was added. Sorting samples were collected on an Aria 561 (BD). Representative plots were made with and quantification was performed by using FlowJo (Tree Star Technologies).

### Library amplification

Genomic DNA was extracted using the NucleoSpin™ Blood kit (Macherey-Nagel, 740951) according to the manufacturer’s instructions. Up to 1 μg of genomic DNA was used for PCR amplification of gRNA sequences for NGS. Amplification was performed using NEBNext Ultra II Q5 Master Mix (New England BioLabs, M0544), with primer sequences listed in table S1. Optimal PCR cycle numbers were determined empirically by amplifying 20 ng of extracted genomic DNA and selecting the lowest number of cycles that produced a visible band on an agarose gel. PCR products were purified using a double-sided SPRI bead size selection (0.65× followed by 1.0×) and eluted in 20 μL of elution buffer. Libraries were sequenced on an Illumina NextSeq platform with a 50-8-8-50 read structure.

### *In vivo* single cell screen (Perturbseq readout)

AAV particles carrying the TF gRNA library for Perturb-Seq were packaged as described in the AAV production and purification section. A single dose of 1.0 × 10^11^ viral genome in a total volume of 50 μl was administered intratracheally. Two weeks after AAV gRNA library delivery, mice were infected intranasally with IAV (described above). Lungs were harvested at the time points indicated in the corresponding experimental schematic, stained and transduced epithelial cells (live, singlet CD45^-^CD31^-^Pdgfra^-^EpCAM^+^mScarlet^+^) were collected. Samples were processed using the 10x Genomics 5’ GEM-X single-cell platform to enable simultaneous transcriptome and gRNA capture. Libraries for gene expression and gRNA identity were constructed and sequenced following the manufacturer’s protocols.

### Evaluation of *Sftpc* and *Msl1* gRNA genome editing efficiency *In vivo*

Single gRNAs were cloned into SAGE and packaged into AAV as described above. To induce Cas9 expression, two doses of tamoxifen were IP administered seven days prior to AAV delivery. For *Sftpc* targeting, SAGE carried the gRNA sequence 5′-CTACAATCACCACGACAACG-3′. For *Msl1*, SAGE carrying two different Msl1 gRNAs were independently packaged and mixed at equal ratio #1 (5′-GCGATAGAGCCCGTCTGAAT) and #2 (5′-GGTCCGAGAGAGATACGGTA). SAGE was administered intratracheally at a dose of 1.0 × 10^11^ viral genomes per mouse. Lungs were harvested 14 days after administration. For the *Sftpc* knockout experiments, the left lobe was fixed and embedded in OCT for sectioning and immunofluorescence, while the right lobes were dissociated as described above to isolate mScarlet^+^eGFP^+^ AT2 cells. For *Msl1* gRNA testing, whole lungs were dissociated for sorting of mScarlet^+^eGFP^+^ AT2 cells. Genomic DNA was extracted from sorted AT2 cells by incubating them in 0.25x CutSmart Buffer (New England BioLabs, B6004) containing 0.4 mg/mL Proteinase K (Qiagen, 19133). Cells were resuspended at 100 cells/µL, digested at 56 °C for 20 min, and then heated at 95 °C for 10 min to inactivate enzymes. 5-10 µL of this lysate was used for PCR amplification with primers flanking the predicted gRNA cut sites (see table S1), using PrimeSTAR GXL polymerase (Takara Bio, R051A) according to the manufacturer’s instructions. PCR products were purified using the NucleoSpin Gel and PCR Clean-up Kit (Macherey-Nagel, 740609) and submitted to Genewiz for Sanger sequencing. Genome editing efficiency was quantified using the ICE analysis tool (Synthego, https://ice.editco.bio/).

## Data analysis

### *In vivo* pooled genetic screen analysis

Raw FASTQ files were processed using the MAGeCK package^92^. The count function was first applied to generate an gRNA read count table. Samples whose gRNA abundance showed a Pearson correlation coefficient below 0.8 with all other samples were excluded from downstream analyses. Subsequently, the test function was used to estimate fold changes and assess the statistical significance of individual gRNAs between experimental conditions.

### *In vivo* perturbSeq analysis

#### scRNA-seq data processing

BCL files of transcriptome libraries were used to generate FASTQ files using the default parameters by ‘cellranger mkfastq’ command (Cell Ranger v8.0.1). The ‘cellranger count’ command was used to align the transcriptome reads to the mouse genome reference mm10 (GENCODE vM23/Ensembl 98) and generate a gene expression count matrix.

#### gRNA data processing

gRNA libraries were demultiplexed into FASTQ files using *bcl2fastq.* Cell barcode, unique molecular identifier (UMI), and gRNA sequences were extracted and summarized from the raw FASTQ files. Cell barcodes with sequencing depth below the 10th percentile were removed. For each remaining cell barcode, the most abundant (top) and second most abundant gRNA sequences were identified and mapped to the gene expression library. To minimize potential gRNA misassignment, only gRNA meeting the following criteria were retained: (i) the top gRNA UMI counts was greater than or equal to the 10th percentile of all top gRNA UMI counts, and (ii) the top gRNA abundance was at least two-fold higher than that of the second abundant gRNA. The top gRNA passing these filters was assigned as the gRNA identities for downstream analysis.

### Cell type classification and cell identity annotation

#### MSL and NSL complex analysis

Filtered count matrices generated by Cell Ranger were imported into R (v4.3.0) using the ReadMtx function from Seurat (v5.0.0) and used to construct Seurat objects with CreateSeuratObject. Cells with fewer than 200 detected genes or with >10% mitochondrial transcript content were excluded. The dataset was log-normalized, and the 2,000 most variable genes were identified using FindVariableFeatures. The data were then scaled with ScaleData, followed by principal component analysis (PCA) using RunPCA. Dimensionality reduction was performed using Uniform Manifold Approximation and Projection (UMAP) via RunUMAP. Clustering was carried out using FindNeighbors and FindClusters on the first 30 principal components with a resolution of 0.8. Cluster identities were manually annotated based on canonical marker gene expression. Clusters identified as low-quality, defined by low gene and UMI counts, were removed from further analysis.

#### Transcription factor screen analysis

For the TF screen, data from each time point was processed independently, using the ReadMtx function from Seurat (v5.0.0) and used to construct Seurat objects with CreateSeuratObject. Cells were filtered by excluding those with <200 genes, >99.9th percentile of gene or UMI counts, or >20% mitochondrial content. Filtered Seurat objects were merged across time points. The merged dataset was log-normalized and the top 2,000 variable features were identified with FindVariableFeatures. Data integration and scaling were performed using ScaleData, followed by PCA with RunPCA. Batch correction across time points was performed using Harmony on the first 30 PCs. The corrected PCs were used for UMAP visualization and clustering via RunUMAP, FindNeighbors, and FindClusters (dims = 1: 30, resolution = 0.8). Cluster identities were annotated using known lineage markers. Non-alveolar epithelial populations, including ciliated cells, club cells, and AT2-club intermediates, were excluded from downstream analysis to focus on alveolar lineage biology.

#### HOTSPOT analysis

We applied HOTSPOT (v1.1.3) to identify statistically significant gene co-expression patterns across the single-cell transcriptomic dataset. Precomputed Harmony embeddings were used as the input representation to account for batch effects. As an initial step, the top 2,500 genes showing significant autocorrelation in expression were selected for module detection. HOTSPOT then clustered these genes into co-expression modules that exhibited coordinated expression patterns across the Harmony-defined transcriptional manifold. Only modules containing more than 30 genes were retained for downstream analysis to ensure robustness and biological interpretability. Each retained gene program was manually annotated based on known functional categories, enriched pathways, or characteristic marker genes. To quantify program activity at the single-cell level, we computed a program score for each cell as the first principal component of the standardized expression matrix restricted to genes within the program subspace. Finally, to assess perturbation effects, program scores were averaged across all cells belonging to the same perturbation condition. These mean program scores were ranked to evaluate and compare the transcriptional impact of each gene knockout on the identified co-expression programs.

#### Transcription factor knockout effects group assignment

To systematically classify transcription factor function, we selected HOTSPOT-derived gene programs representing key epithelial states or biological processes including AT2, AT1, DATP, aberrant, inflammation, stress, and proliferation programs. TFs associated with fewer than 100 cells were excluded. For each TF, the mean knockout-induced change in program score was computed by centering per-cell program scores to the control population.

TFs were first grouped based on their effects on the AT2 program. TF knockouts with average AT2 program scores in the top 25% quantile were classified as AT2 retention. Among TFs that reduced or minimally altered AT2 identity, we computed Mahalanobis-standardized contrasts between AT1 and aberrant program deltas, along with overall terminalization magnitude (sum of positive AT1 and aberrant deltas) and co-activation level (ratio of the smaller to the total positive component). TF knockout effects were then categorized into AT1-biased, aberrant-biased, or mixed terminalizing groups using data-adaptive thresholds, defined as the 80th percentile of terminalization magnitude and the 70th percentiles of bias and mixedness metrics among strong terminalizers. TFs with weak or negligible effects on terminal programs were labeled as low-impact, and threshold-based fallback rules were applied to resolve borderline cases. Knockout effects of *Hmgb1* and *Carhsp1* are not included due to low cell numbers (<100).

#### Transcription factor co-binding analysis

To generate ATAC-seq peak regions for transcription factor co-binding analysis, BigWig signal files from Choi et al., 2020 were processed using R packages rtracklayer (v1.62.0), GenomicRanges (v1.52.1) and data.table (v1.14.10). First, signal values were extracted, and peaks were defined by applying a quantile-based threshold; after testing multiple cutoffs (0.90, 0.85, and 0.80), regions above the 80th percentile of signal intensity were retained. Adjacent high-signal regions within 100 bp were merged to define continuous peaks. The resulting genomic intervals were exported as BED files and used for downstream motif and co-binding analyses with HOMER.

To assess TF co-binding within promoter regions, we integrated the above ATAC-seq accessibility peak regions with publicly available TF ChIP-seq datasets from ReMap 2022 (https://remap2022.univ-amu.fr/). Gene annotations for protein-coding genes were retrieved from Ensembl (release 102), and TSS were defined using biomaRt package (v2.58.2) and converted into GRanges objects. Promoter regions were defined as −2000 to +500 bp relative to each TSS. ATAC-seq peaks BED files (retained at ≥ 100 bp) were intersected with promoter regions to identify accessible promoter intervals. ChIP-seq binding sites for selected TFs (Etv5, Cebpa, Max, Foxa2) were restricted to standard chromosomes. Overlaps between promoter-associated ATAC-seq peaks and TF binding sites were quantified using *GenomicRanges*, and co-binding scores were computed by summing TF occupancy per gene. Genes of interest (AT1 program gene list from HOTSPOT analysis) were then compared with the background (all genes not in the list) set using Wilcoxon rank-sum tests, and co-binding patterns were visualized using UpSet plots using the ComplexUpset (v1.3.3).

Motif enrichment was then performed using HOMER (v5.1), applying the findMotifsGenome.pl command with the -size given option. Selected enrichments for known motifs were reported.

#### Comparing DATP-like states with human PF data

Human pulmonary fibrosis single-cell datasets from Adams et al., 2020 (GSE136831) and Habermann et al., 2020 (GSE135893) were obtained from NCBI GEO. For Adams et al., epithelial cells were subset, log-normalized, and the top 2,000 variable features identified using *FindVariableFeatures*. Data were scaled (*ScaleData*), reduced via PCA (*RunPCA*), and visualized using UMAP (*RunUMAP*). Clustering was performed with *FindNeighbors* and *FindClusters* (dims = 1:40, resolution = 0.8). Original cell type annotations were retained.

For Habermann et al., epithelial cells were subset using original annotations. The data were log-normalized, variable features selected, and batch correction across time points applied using *Harmony* on the top 30 PCs. UMAP and clustering (*FindNeighbors*, *FindClusters*) were performed using the corrected PCs (dims = 1:30, resolution = 1.0). Original cell type labels were preserved.

Gene sets from Wang et al., 2023 representing senescent, AT2, AT2 transitional, and basaloid states were used to compare expression profiles between AT2 and DATP-like cells in mouse dataset and AT2, transitional AT2, and basaloid cells in human dataset.

#### Trajectory analysis

To reconstruct differentiation trajectories of alveolar epithelial cells, we performed pseudotime analysis using diffusion maps coupled with Slingshot (v2.8.0). Cells annotated as AT2, AT2-AT1, AT1, DATP-like-1, DATP-like-2, and aberrant were subset from the single-cell dataset while retaining Harmony-integrated principal components. A stratified subset of 10,000 cells balanced across cell states and timepoints was used to train the diffusion map in order to reduce computational complexity while maintaining representative coverage. Diffusion components were computed with the destiny package (v3.16.0, k = 60, local sigma), and all cells were projected into the trained diffusion map space using k-nearest neighbor averaging.

Lineage inference was then performed using the Slingshot package with diffusion map embeddings as input, specifying AT2 as the root cluster and AT1 and aberrant states as terminal endpoints. Slingshot provided lineage-specific pseudotime estimates and curve weights, which were used to quantify differentiation bias toward AT1 versus aberrant outcomes. Branch bias was defined as the difference in Slingshot-assigned curve weights between AT1-directed and aberrant-directed lineages for each cell.

Visualization was carried out using the ggplot2 package (v3.4.4). Diffusion components DC1 and DC2 were plotted with cells colored by state or branch bias, and Slingshot-inferred curves were overlaid.

#### Pathway activity scoring

Pathway activity was assessed using curated human and mouse gene sets from HALLMARK_INTERFERON_GAMMA_RESPONSE, *HALLMARK_MYC_TARGETS_V1*, *HALLMARK_IL6_JAK_STAT3_SIGNALING*, and *HALLMARK_TNFA_SIGNALING_VIA_NFKB* to evaluate NF-*κ*B and Myc signaling activity, respectively. UCell (v2.6.2) scoring was applied to compute enrichment scores for each cell. Scores were compared between DATP-like-1 and DATP-like-2 states, as well as between basaloid and transitional AT2 states, using Wilcoxon rank-sum tests.

---

**Fig. S1.**
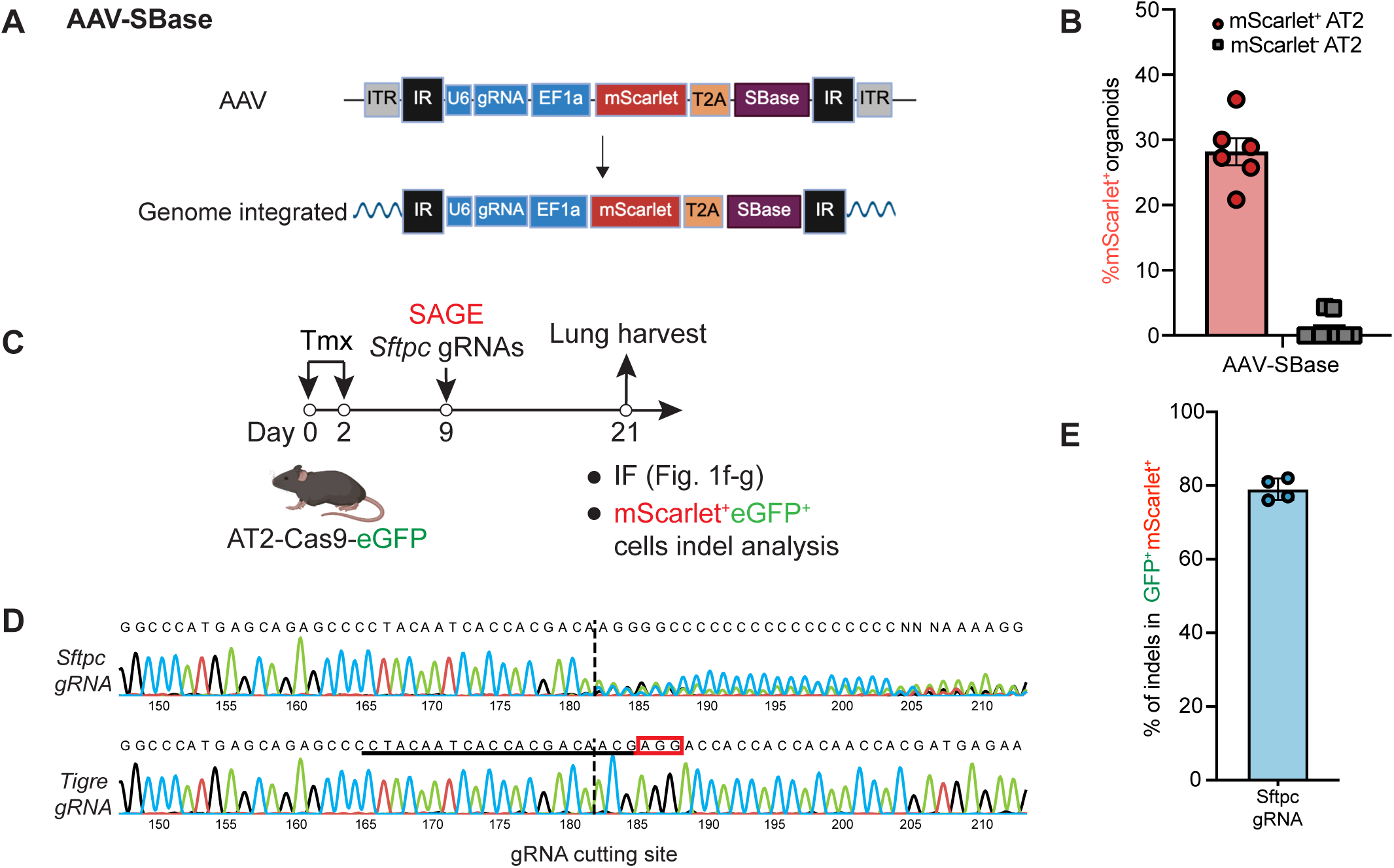
Design and validation of the engineered AAV systems for *in vivo* genome editing. **(A).** Schematic of the AAV-SBase system. **(B).** Quantification for integration efficiency of the AAV-SBase system using the AT2 clonal organoid assay. Error bars: mean ± SEM. **(C).** Schematic workflow for validating SAGE genome editing activity *in vivo*. Tmx, tamoxifen. IF, immunofluorescence. **(D).** Representative Sanger sequencing traces for *Tigre* gRNA (control) and *Sftpc* gRNA near gRNA binding sites. Insertions and deletions (indels) were generated specifically when *Sftpc* targeting gRNA is used. Red square indicates the PAM sequence. **(E).** Indel efficiency for *Sftpc* knockouts. N = 4 mice. Error bars: mean ± SEM.

**Fig. S2.**
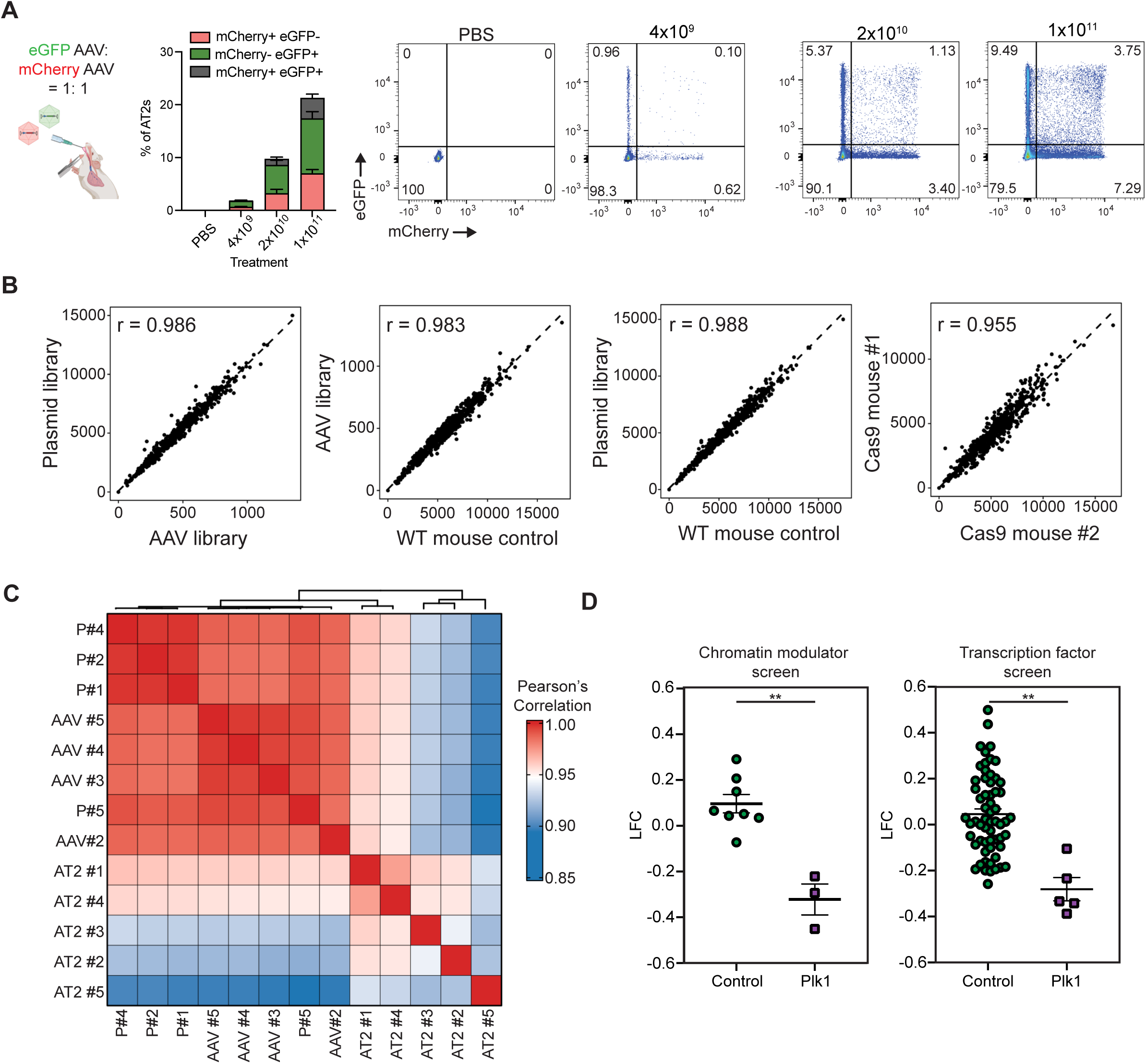
Validation of the SAGE platform for robust library delivery and screening. **(A).** Two color AAV experiments for quantitative assessment of multi-infection rate for AAV transduction *in vivo*. Mice were i.t. administered non-engineered eGFP-AAV9: mCherry-AAV9 at 1:1 ratio at indicated total viral loads (4 × 10^9^, 2 × 10^10^, or 1 × 10^11^) and lungs were harvested five days later. Representative gates were pregated as CD45^-^CD31^-^Pdgfra^-^EpCAM^+^CD104^-^MHCII^+^. **(B).** Pairwise scatter plots of gRNA abundance in libraries from plasmids, AAVs, and AT2s from biological replicates. **(C).** Pearson correlation of gRNA abundance in the CM screen derived from plasmids (P), packaged AAV libraries (AAV), and AT2s harvested after injury resolution. Pairwise correlations demonstrate high library representation fidelity across stages. **(D).** gRNA abundance log_2_ fold change (LFC) of control gRNAs and *Plk1* gRNAs in the CM and TF screens. Error bars: mean ± SEM. Statistical analysis was using the two-tailed Student’s *t*-tests. p-values are reported as follows: **p < 0.01.

**Fig. S3.**
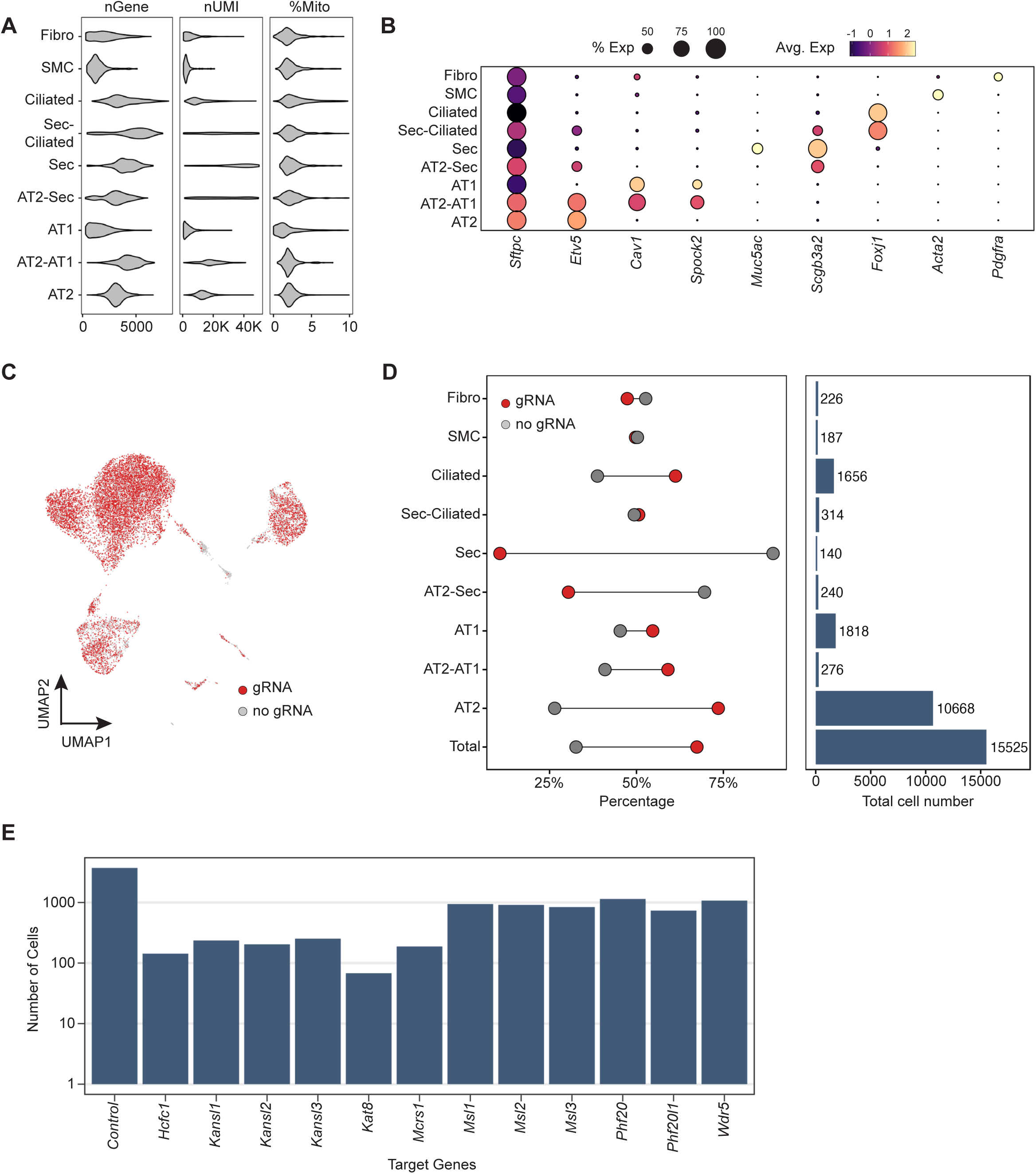
Quality control and gRNA capture metrics for SAGE-Perturb-seq. **(A).** Violin plots of quality control metrics for number of detected genes (nGenes), total UMI counts (nUMI), and percentage of mitochondrial reads (%Mito) across cell populations. UMI, unique molecular identifiers. **(B).** Representative marker gene expression across cell populations. **(C).** UMAP representation of the distribution of cells with and without gRNA assignment. **(D).** gRNA capture rate distribution across cell populations. **(E).** Distribution of the number of cells with gRNA assignment for each gene target.

**Fig. S4.**
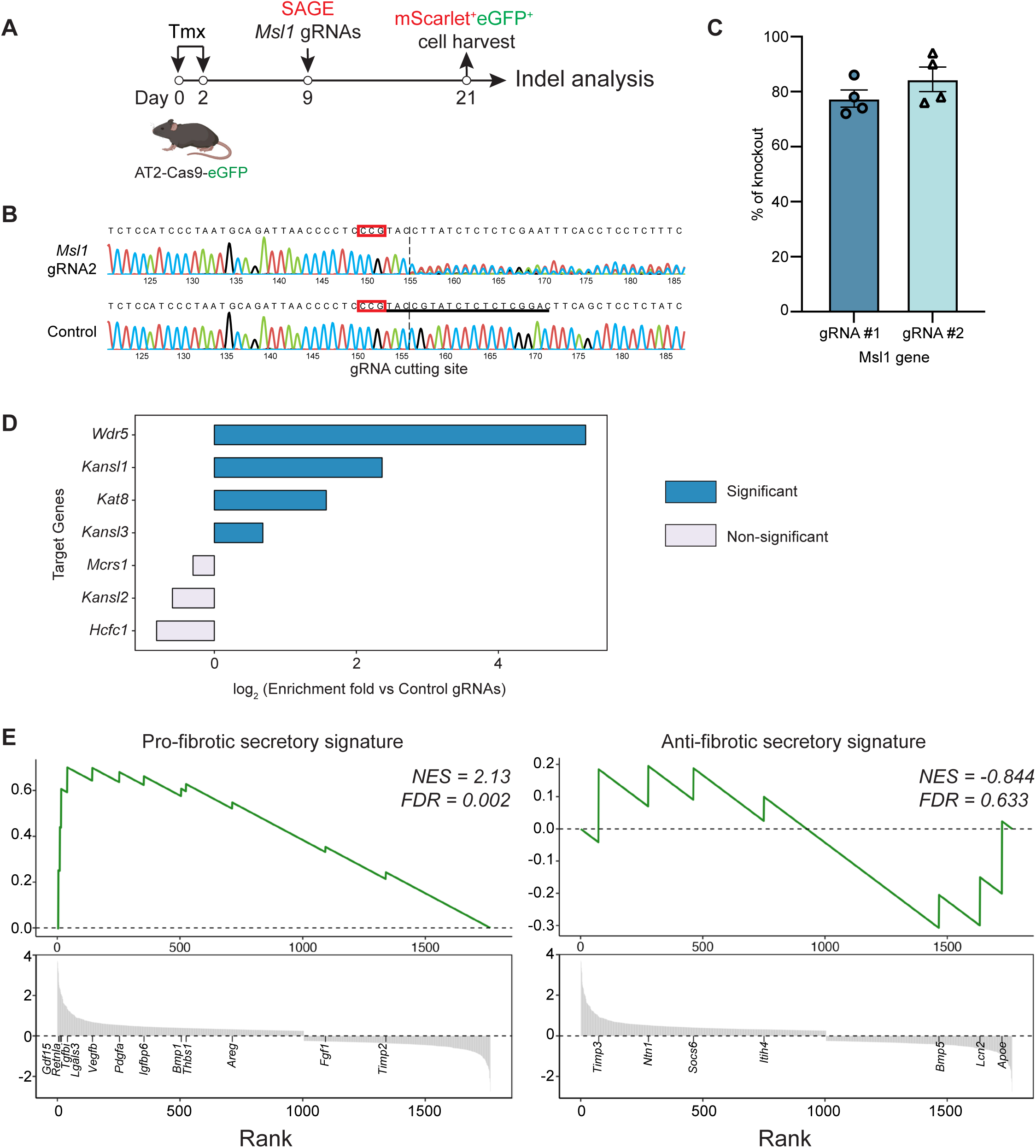
Validation of NSL complex perturbations and functional consequences in AT2s. **(A).** Schematic workflow for quantification of *Msl1* gRNA genome editing efficiency *in vivo*. **(B).** Representative Sanger sequencing traces for control gRNA (*Tigre*) and *Msl1* gRNA #2 at *Msl1* gene locus. Indels were generated specifically when *Msl1* targeting gRNAs were used. Red squares indicate the reverse complementary sequence for PAM. **(C).** Quantification of indel efficiency of *Msl1* gRNAs. N = 4 mice. Error bars: mean ± SEM. **(D).** Enrichment of gRNAs targeting individual NSL essential components in comparison to control gRNAs. Fisher’s exact test was used. p-values were adjusted for multiple comparisons using the Benjamini-Hochberg approach to control for false discovery rate. Significance was defined as FDR < 0.1, and corresponding bars are highlighted in blue. **(E).** AT2-2 upregulated profibrotic gene signatures. Gene set enrichment analysis (GESA) was performed using differentially expressed genes between AT2-2 and AT2-1 subpopulations. NES, normalized enrichment score.

**Fig. S5.**
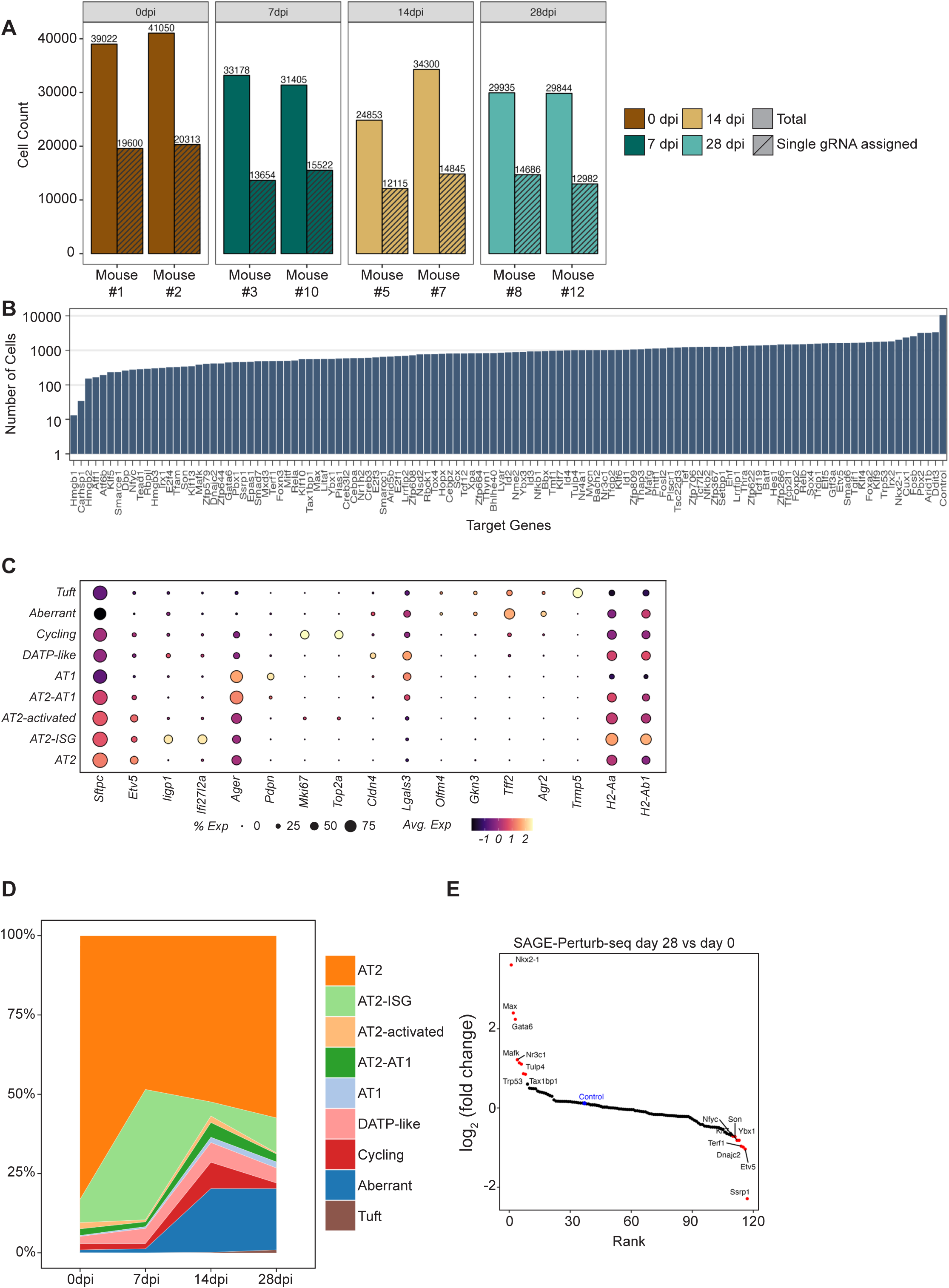
Overview and quality metrics of the SAGE-Perturb-seq atlas on transcription factor knockouts. **(A).** Cell recovery and single gRNA assignments across time points. **(B).** Distribution of cell numbers across different gRNA gene targets. **(C).** Representative marker gene expression for different cell populations in SAGE-Perturb-seq TF knockout dataset. **(D).** Dynamic changes of cell proportions during IAV induced injury and repair. **(E).** Comparison of gRNA abundance between the day 28 and day 0 time points. Each point represents an individual gRNA, ranked by log_2_ fold change. gRNAs with high log_2_ fold change >1 or low log_2_ fold change < −1 are highlighted in red. Control gRNAs are grouped, averaged together, and labeled in blue.

**Fig. S6.**
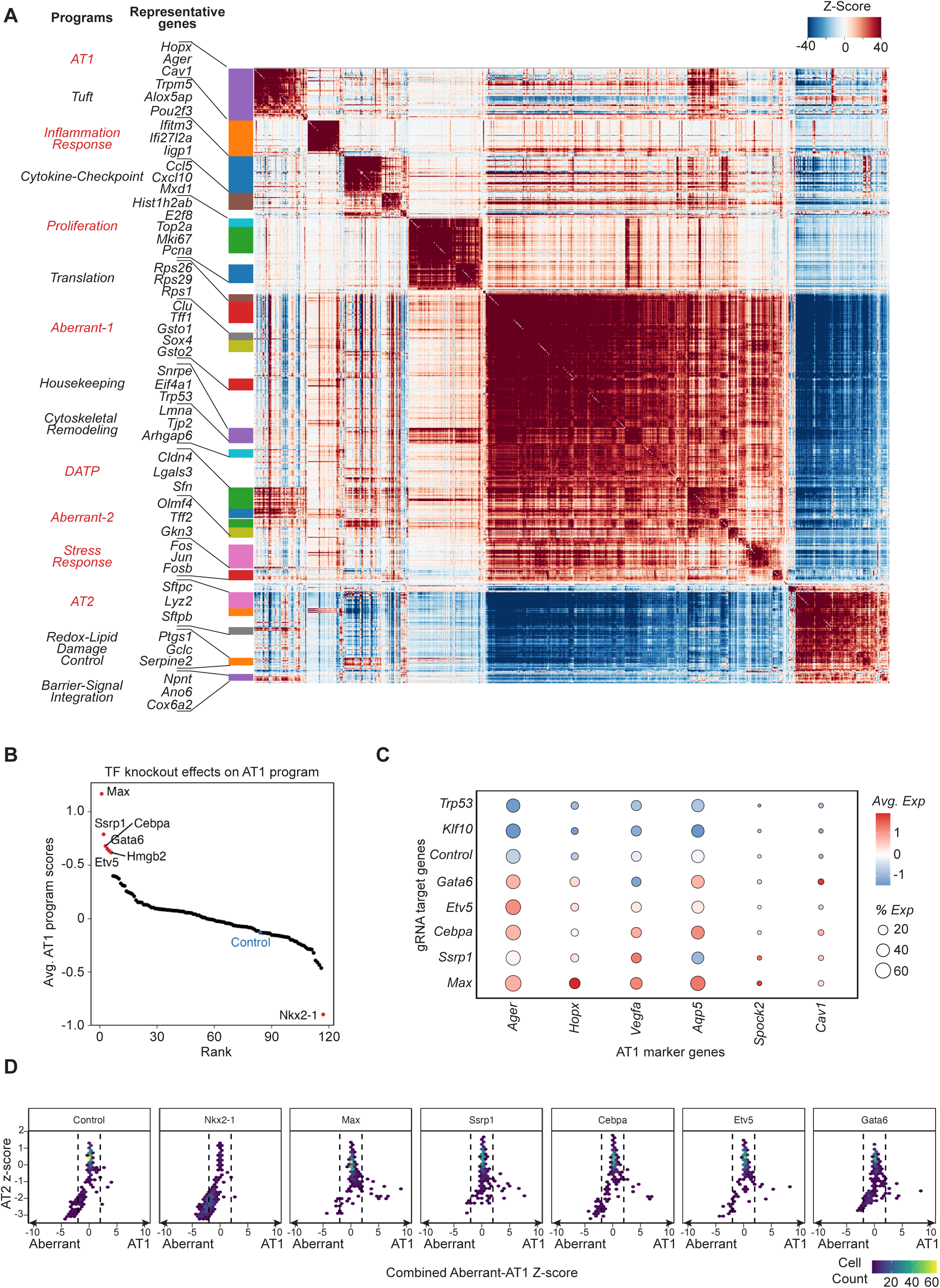
Hotspot analysis and functional mapping of transcription factor perturbations influencing AT1 programs. **(A).** Hotspot analysis of gene expression programs. Hotspot identified 25 gene programs based on pairwise local correlations of gene expression. Similar gene programs are grouped together and consolidated into 14 programs. Program names and representative genes are listed on the left. Gene programs selected for TF knockout effect grouping are highlighted in red. **(B).** Rank plot of average AT1 program scores per gRNA target. gRNAs with high (avg. score > 0.5) or low (avg. score < −0.5) AT1 program scores are highlighted in red. Control gRNAs are grouped, averaged together, and labeled in blue. **(C).** Expression of representative AT1 marker genes in cells with selected gene knockouts. **(D).** Hexbin plots visualization of AT2 program activity relative to lineage differentiation across selected gene knockouts. The x-axis represents the first principal component (PC1) derived from AT1 and dysplastic program scores, and the y-axis indicates the z-scored AT2 program activity. Color intensity reflects cell density.

**Fig. S7.**
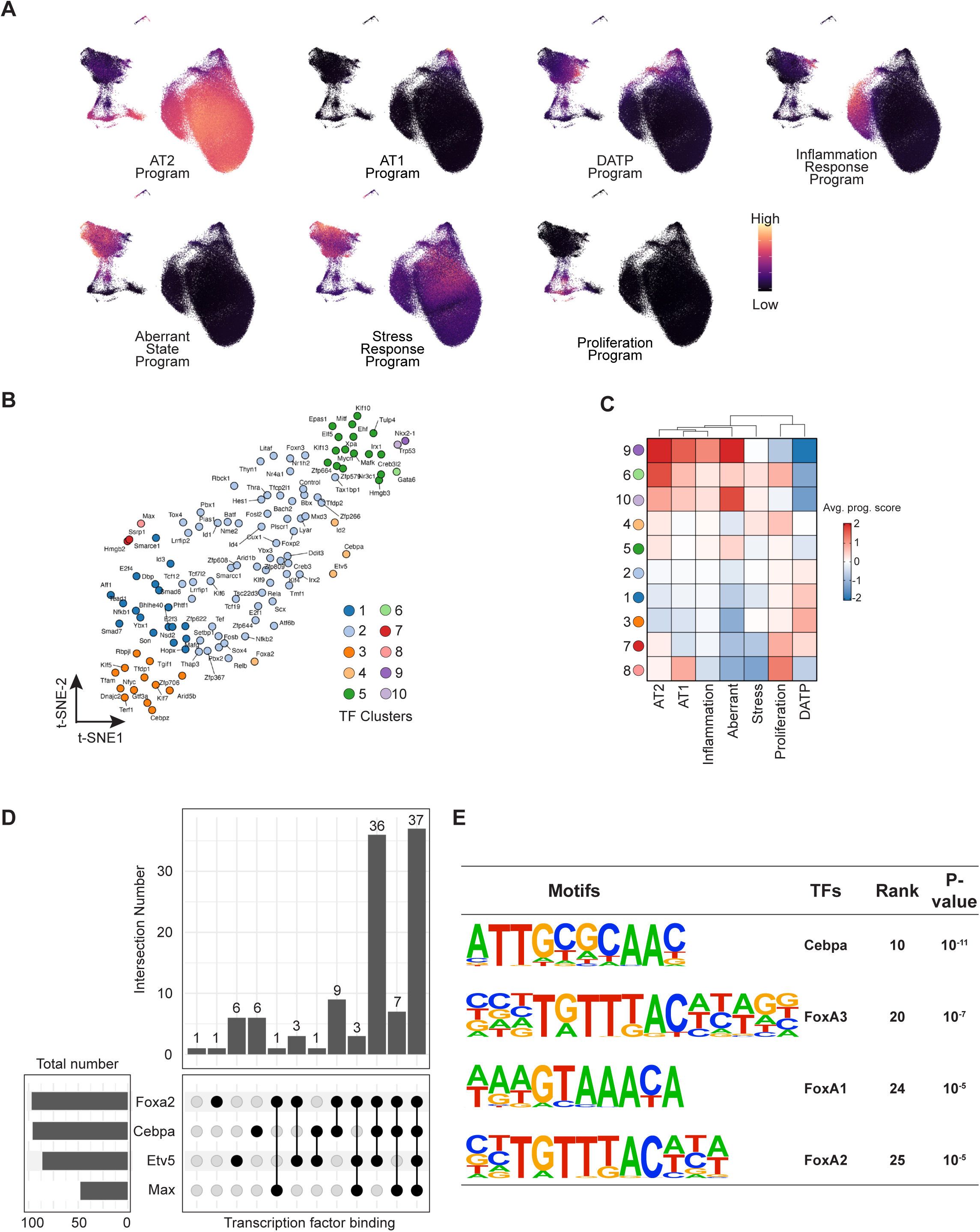
Functional mapping of transcription factor perturbation effects in alveolar repair. **(A).** UMAP representation of selected gene program enrichment. **(B).** Display of knockout effects on the t-SNE map. Each point represents an individual TF knockout. Average program scores from selected gene programs were used as input for K-means clustering. Colors indicate clusters of knockouts with similar functional impact. **(C).** Heatmap of average program scores across TF knockout clusters. For each cluster, Average program scores were computed and visualized, with values capped between −2 and 2. **(D).** UpSet plot of TF co-binding on AT1 program gene promoters. Publicly available ChIP-seq datasets (ReMap2022) for Foxa2, Cebpa, Etv5, and Max were used to define co-binding events. Promoter regions were defined as −2000 to +500 bp around the TSS and restricted to loci overlapping with ATAC-seq open chromatin regions in AT2 cells. **(E).** HOMER known motif enrichment analysis of AT1 program-associated regulatory regions. Known motif enrichments were identified using HOMER from ATAC-seq open chromatin regions in AT2s linked to AT1 program genes.

**Fig. S8.**
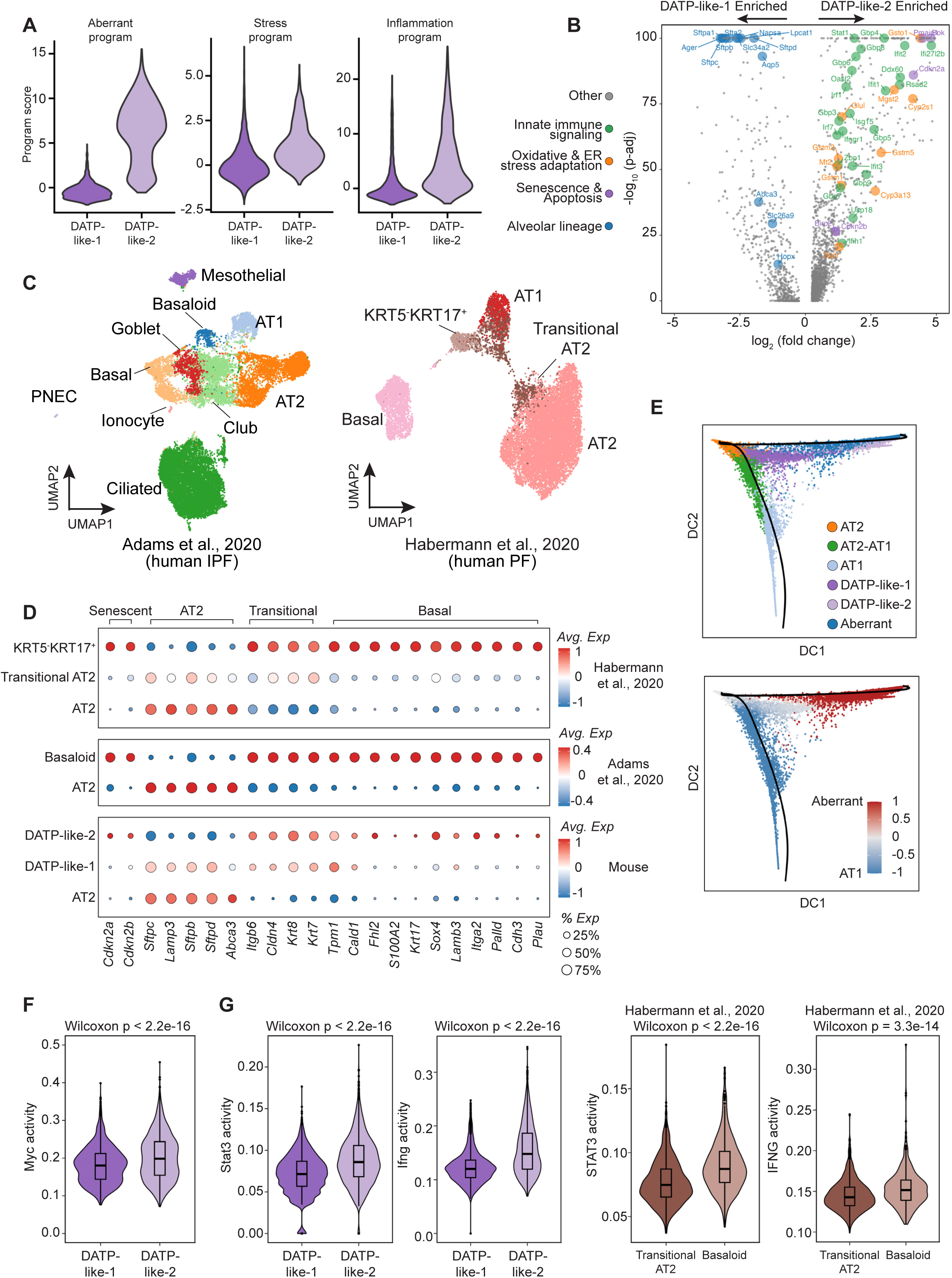
DATP-like-2 state exhibits features of human basaloid cells. **(A).** Violin plots of selected program score distributions in DATP-like-1 and DATP-like-2 states. **(B).** Volcano plots depicting differentially expressed genes between DATP-like-1 and DATP-like-2 states. Significantly differentially expressed genes are defined by log_2_(fold change) > 1 or <-1 and adjusted p value < 0.05. Selected functional groups are color-highlighted. **(C).** UMAP representation of major epithelial cell populations from publicly available human PF single cell RNA-Seq data^19,21^. **(D).** Comparison of mouse and human transitional states. Gene list was selected from Wang et al., 2023. **(E).** Diffusion map illustrating distinct fate trajectories adopted by transitional cells. The top panel is color-coded by cell populations, and the bottom panel highlights the corresponding fate biases. **(F).** Myc activity in DATP-like states. Statistical significance was assessed using the Wilcoxon rank-sum test. The center lines indicate the medians, and the boxes represent the interquartile ranges. **(G).** Inflammatory pathway activity (Stat3 and Ifng) in pathological transitional states in both mouse and human PF lungs^21^. UCell^94^ scoring was applied to compute enrichment scores for each cell. Statistical significance was assessed using the Wilcoxon rank-sum test. The center lines indicate the medians, and the boxes represent the interquartile ranges.

